# Networks of descending neurons transform command-like signals into population-based behavioral control

**DOI:** 10.1101/2023.09.11.557103

**Authors:** Jonas Braun, Femke Hurtak, Sibo Wang-Chen, Pavan Ramdya

## Abstract

To transform intentions into actions, movement instructions must pass from the brain to downstream motor circuits through descending neurons (DNs). These include small sets of command-like neurons that are sufficient to drive behaviors—the circuit mechanisms for which remain unclear. Here, we show that command-like DNs in *Drosophila* directly recruit networks of additional DNs to orchestrate flexible behaviors. Specifically, we found that optogenetic activation of command-like DNs previously thought to drive behaviors alone in fact co-activate larger populations of DNs. Connectome analysis revealed that this functional recruitment can be explained by direct excitatory connections between command-like DNs and networks of interconnected DNs in the brain. The size of downstream DN networks is predictive of whether descending population recruitment is necessary to generate a complete behavior: DNs with many downstream descending partners require network recruitment to drive flexible behaviors, while neurons with fewer partners can alone drive stereotyped behaviors and simple movements. Finally, DN networks reside within behavior-specific clusters that inhibit one another. These results support a mechanism for command-like descending control whereby a continuum of stereotyped to flexible behaviors are generated through the recruitment of increasingly large DN networks which likely construct a complete behavior by combining multiple motor subroutines.

## Introduction

Animals, including humans, are capable of generating a remarkable variety of behaviors. For example, they can navigate rugged terrain and manipulate objects in their environments by coordinating their limbs. These movements ultimately arise from motor circuit dynamics in the vertebrate spinal cord or invertebrate ventral nerve cord. Nevertheless, the selection, initiation, and control of behaviors often depends on ongoing commands sent from the brain to downstream motor circuits via a relatively small population of brain neurons called descending neurons (DNs).

We still lack mechanistic understanding of how DNs as a population drive and coordinate flexible behaviors that require ongoing feedback (e.g., goal-directed walking or reaching for an object) as well as those which are more stereotyped (e.g., escape). This is in part due to the technical difficulty of comprehensively recording and manipulating DNs in behaving mammals: there are *>*1 million in the human pyramidal tract ^1^ and *∼*70,000 in the mouse corticospinal tract ^2^. By contrast, the adult fly, *Drosophila melanogaster*, has approximately 1,300 DNs linking the brain to motor centers in the ventral nerve cord (VNC) ^3^. Despite this numerical simplicity, flies can generate a variety of complex behaviors including walking over challenging terrain ^4^, flight ^5^, courtship ^6^, and aggression ^7^. Recently, it has become possible to quantify the synaptic connectivity of every neuron—including DNs—in the brain ^3^ and the VNC ^8,9^. Importantly, in flies DNs can be repeatedly targeted across individual animals for experimental recordings (electrophysiological ^10^ or optical ^11^) or manipulations (e.g., activation ^12^ or silencing ^13^) using genetic lines that drive transgene expression in small sets of DNs ^14,15^.

One notable discovery that was derived using these tools was the observation that, despite the abundance of DNs in the fly brain, artificial activation of very small sets (2-4 neurons) of so-called ‘command-like’ DNs can be sufficient to drive a complete behavior (but not strictly necessary as for ‘command’ neurons ^16^). For example, DNs have been identified whose artificial activation triggers forward walking (DNp09) ^17^, anterior grooming (antennal DNs / aDN, DNg11, DNg12) ^18,19^, backward walking (moonwalker DNs / MDN) ^20^, escape (giant fiber neurons / GF) ^10^, egg-laying (oviDN) ^21^, and aspects of courtship (pIP10, aSP22) ^22,23^. The capacity of some DNs to act as command-like neurons appears to be general across species including other invertebrates (e.g., crayfish giant escape neurons ^24^ and cricket song neurons ^25^) and mammals (e.g., neurons which halt locomotion in mice ^26^). This framework of command-like descending control—using simple, low-dimensional brain signals to drive downstream distributed motor circuits—has been formalized into controllers for bio-inspired robots that can walk and swim ^27^.

The concept of command-like control raises a fundamental question: To what extent does every pair or small set of DNs drive a distinct behavior? Several lines of evidence refute this possibility. Most directly, for many DNs, sparse optogenetic activation does not clearly and reliably drive a distinct, coordinated behavior ^12^. Additionally, even during one behavior—forward walking and turning—we have observed that populations of many DNs in the cerebral ganglia become co-active ^28^. This is in line with the observation that a population of 15 DNs can modulate wing beat amplitude ^29^ and that the activation of individual DNs has a lower probability of driving take-off than co-activation of multiple DNs ^30^. Beyond controlling kinematics, it has also been shown that DNs can convey sensory information ^28,31^ and that they may be modulatory ^32,33,34,35^. All of these observations suggest that, rather than being low-dimensional, DN control of behavior is population-based or high-dimensional: the brain flexibly engages larger populations of DNs to construct and mediate complete behaviors.

At first glance, these two models—command-like versus population-based DN behavioral control— appear to be incompatible. However, we can envision at least two means of unifying these seemingly disparate observations. On one hand, command-like or non-command-like DNs may simply target different downstream motor circuits (i.e., in the spinal cord or VNC) that can or cannot generate complete behaviors, respectively. On the other hand, command-like DNs might be privileged in that they can recruit additional DN populations to drive complete behaviors. This latter possibility is supported by the fact that, in addition to projecting to the VNC, 85 % of all DNs have axon collaterals in the brain’s gnathal ganglia (GNG), where the majority of DNs also originate ^14^. Thus, the GNG may function as a locus where command-like DNs engage other DNs to construct complex behaviors from simpler motor primitives.

Here we investigated the degree to which command-like DNs and other DNs interact with one another in the brain to drive complete behaviors. While optogenetically activating three sets of command-like DNs driving a range of distinct actions, we observed the co-activation of additional DN populations in the GNG. The degree of this functional recruitment covaries with and can be explained by excitatory monosynaptic connections between command-like DNs and downstream DN networks. Through decapitation experiments, we show that this engagement of DN networks is necessary to drive complete flexible behaviors like forward walking and grooming but not for stereotyped behaviors like backward walking. This predicts that command-like DNs driving larger DN networks require their downstream partners in the brain to execute a behavior while DNs with smaller networks do not—a prediction we validate for six additional sets of DNs. Finally, a comprehensive analysis of all DN-to-DN interconnectivity in the brain revealed that DN networks form clusters that are linked to distinct behaviors and that largely inhibit one another. These findings can reconcile the two dominant models of DN control: Command-like DNs drive behaviors by recruiting additional DN populations— the extent to which depends on the flexibility versus stereotypy of the behavior in question. These DN populations likely construct complete behaviors by combining multiple DN-driven motor subroutines.

## Results

### An optogenetic approach to investigate the relationship between command-like DNs and DN population activity

We set out to explore the relationship between two prominent models for how DNs control behavior. In the first model, the artificial activation of a few ‘command-like’ DNs (‘comDNs’)—a simple high-level descending signal—engages downstream motor circuits in the VNC and is thus sufficient to drive a complete behavior (e.g., walking or grooming) **(Fig. 1a, left ‘comDNs’)**. In the second model, a larger population of DNs must become co-active to drive a given behavior. Each DN within this population would be responsible for controlling or modulating a particular motor primitive. The combined activity of the entire population would yield a complete behavior **(Fig. 1a, right ‘popDNs’)**.

**Fig. 1:**
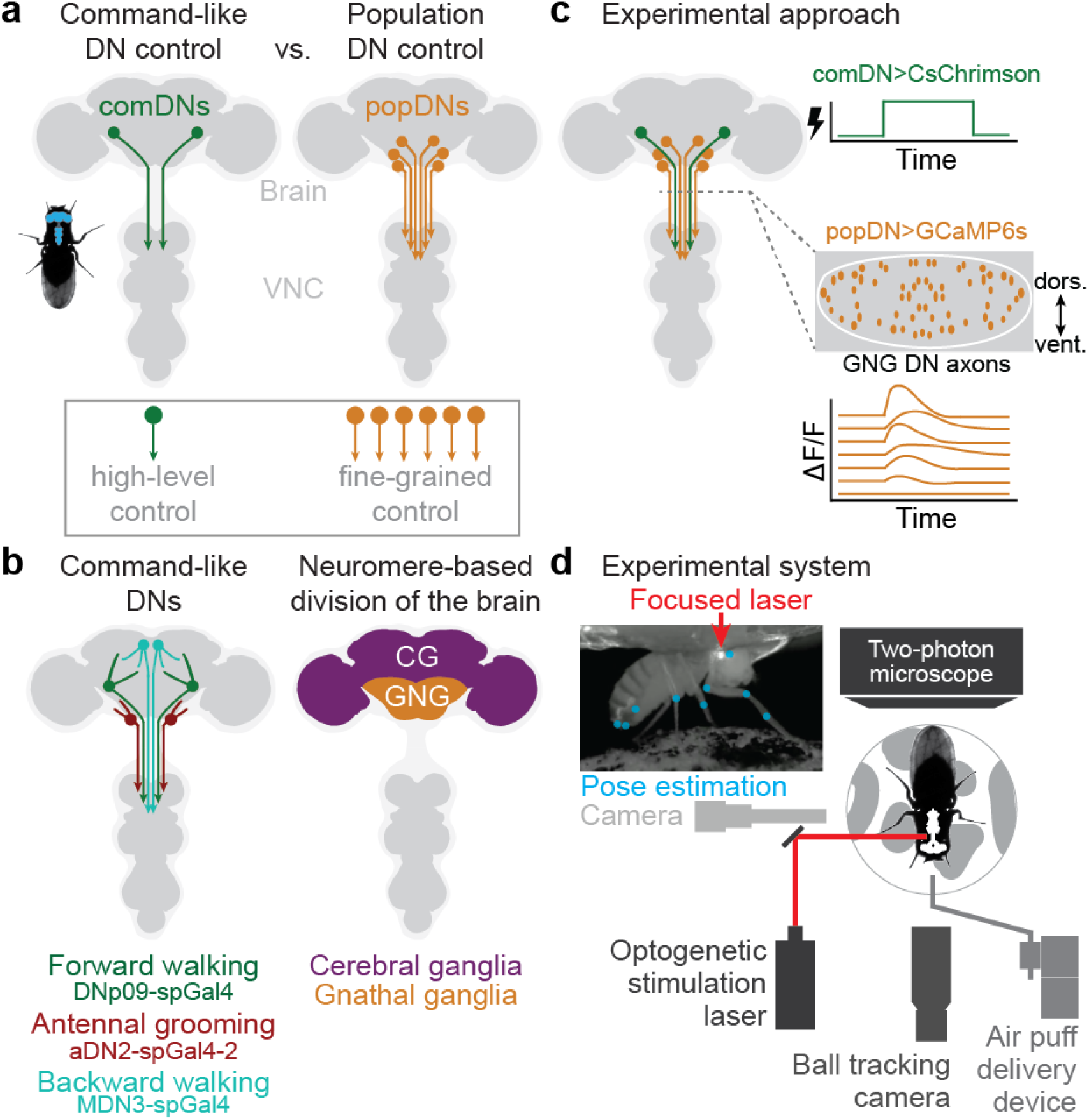
Optical approach to probe the relationship between command-like descending neurons and populations of descending neurons in behaving animals. **(a, left)** Schematic of the *Drosophila* nervous system showing a pair of descending neurons (DNs) which project from the brain to motor circuits in the ventral nerve cord (VNC). Activation of small sets of command-like DNs (‘comDNs’, green) can drive complete actions like walking and grooming. Thus, command-like DNs are thought to send simple, high-level control signals to the VNC, where they are transformed into complex, multi-joint movements. **(a, right)** However, much larger populations of DNs (‘popDNs’, orange) are known to become active during natural walking and grooming. Therefore, in another model, individual DNs contribute to complex behaviors by sending low-level signals to control the fine-grained movements of individual or sparse sets of joints. **(b, left)** A schema illustrating three sets of DNs which exemplify command-like control for limb-dependent behaviors. These elicit (i) forward walking (DNp09, green) ^14,17^, (ii) antennal or anterior grooming (aDN2, red) ^18^, or (iii) backward walking (MDN, cyan) ^20^. Indicated is the approximate location of DN cell bodies within the brain. VNC targets are not schematized. **(b, right)** Two coarse subdivisions of the adult *Drosophila* brain are the cerebral ganglia (CG) and gnathal ganglia (GNG). These are delineated by neuromere boundaries ^74^. The majority of all DNs and their brain targets are predominantly found in the GNG ^14^. Our neural recordings were restricted to DNs within the GNG (GNG-DNs). **(c)** To uncover the relationship between command-like and population-based DN control, we recorded neural activity in the axons of GNG-DN populations (orange) while optogenetically activating different sets of command-like DNs (green). Indicated (dashed gray line) is the coronal imaging region-of-interest in the thoracic cervical connective. Small orange ellipses represent the cross-sections of descending neuron axons. **(d)** To accomplish these experiments, we used a behavior and neural recording system from ^28^ and added an optogenetic stimulation laser (640 nm) that was focused onto DN axons passing through the neck connective. Cartoon schema is not to scale. Inset shows a real camera image of a fly on the spherical treadmill with focused laser light shining on the neck (red arrow). Measured body and leg keypoints for pose estimation are superimposed (light blue).

These two scenarios can be distinguished by the degree to which activation of command-like DNs further activates other downstream DNs. We tested this by devising an all-optical experimental strategy in *Drosophila melanogaster*, where we could optogenetically activate command-like DNs while recording the activity of DN populations within the GNG (GNG-DNs), the most caudal region of the fly brain. We selected GNG-DNs because *∼*60% of all DNs have their cell bodies in the GNG and *∼*85% of all DNs have axonal output in this region ^14^ **(Fig. 1b, right)**. To explore the range of DN control, we performed this experiment across three sets of command-like DNs driving diverse behaviors. First, we stimulated forward walking, a flexible, goal-directed behavior that can employ closed-loop control ^36^ (DNp09-spGAL4, green). Second, we studied anterior grooming, a somewhat flexible behavior that consists of the sequential control of multiple sub-behaviors ^37^ (aDN2-spGAL4, red), and backward walking, a stereotyped behavior that, although kinematically as complex as forward walking, has been shown to arise principally from stereotyped oscillations of the hindlegs ^38^ (MDN3-spGAL4, cyan) **(Fig. 1b, left)**. We artificially activated these command-like DNs via cell-specific expression of the light-activated ion channel, CsChrimson ^39^ (comDN-spGAL4 *>* UAS-CsChrimson; **Extended Data Fig. 1a,d**).

While stimulating these command-like DNs, we recorded neural activity in GNG DN populations by expressing the genetically-encoded calcium indicator GCaMP6s ^40^ in cells positive for the Hox-gene Deformed (Dfd) ^41^ (*Dfd*-LexA *>* LexAOp-opGCaMP6s). This yields selective expression of GCaMP in the GNG **(Extended Data Fig. 1a,b)** and in the proboscis. To further restrict our neural recordings to DNs, we performed two-photon microscopy of DN axons passing through the thoracic cervical connective, as in ref. ^28^ **(Fig. 1c)**. To increase the specificity of command-like DN optogenetic activation, we designed a stimulation system that focuses a 640 nm laser onto the animal’s neck connective **(Fig. 1d, red)**. This reduced the degree to which our laser light might also activate other brain and VNC neurons **(Extended Data Fig. 1e,f)**.

### Activation of command-like DNs recruits additional DN populations

Using these tools we examined whether additional DNs in the GNG might be recruited upon optogenetic activation of command-like DNs. We used an open-loop trial structure in which 5 s periods with optogenetic stimulation of command-like DNs were interleaved with 10 s periods of spontaneous animal behavior. This approach elicited robust behavioral responses which could be trial-averaged for quantification **(Fig. 2a)**. Similar to what we observed when recording from DNs in the cerebral ganglia ^28^, many GNG-DNs were active during spontaneous behavior in the absence of optogenetic stimulation. Thus, to distinguish between GNG-DN activity due to command-like stimulation versus the initiation of spontaneous behaviors, we only analyzed trials for which flies were walking immediately prior to optogenetic stimulation.

**Fig. 2:**
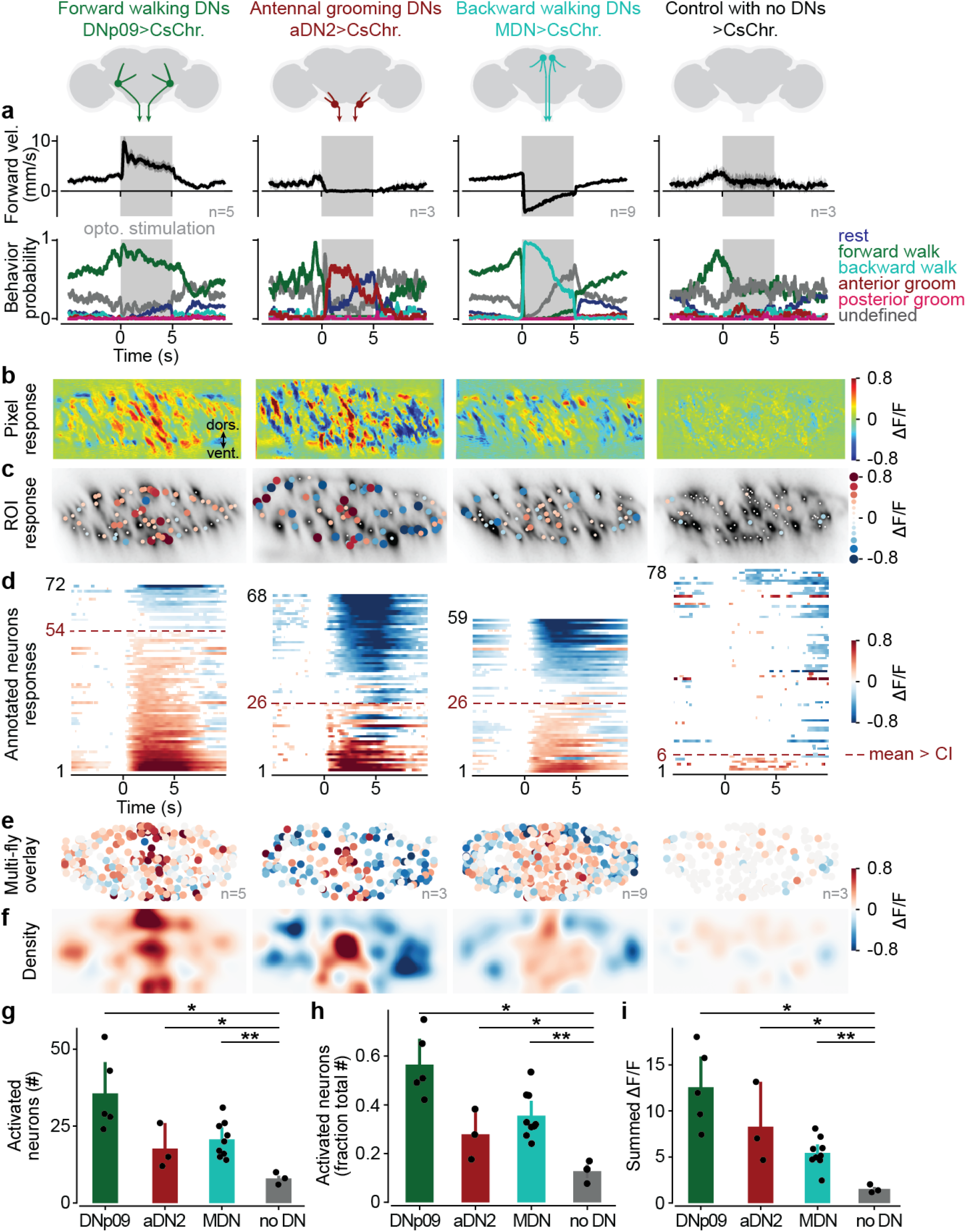
Activation of command-like DNs recruits larger, distinct DN populations. Three command-like DN driver lines were optogenetically stimulated. DNp09 drove forward walking (n=5 flies, total of 120 stimulation trials), aDN2 drove anterior grooming (n=3 flies, total of 34 stimulation trials), and activating MDN drove backward walking (n=9 flies, total of 271 stimulation trials). Control animals without DN expression were also optically stimulated (n=3 flies, total of 47 stimulation trials). **(a)** Shown are forward walking velocities inferred from spherical treadmill rotations (top) and the probability of generating each classified behavior (bottom) during optogenetic stimulation (grey epoch). Shaded area indicates 95 % confidence interval of the mean across all trials. **(b)** Processed two-photon microscopy images illustrating the activation of GNG-DN populations upon command-like DN stimulation. One example animal is shown per driver line (the flies and color scheme are the same as in **Supplementary Videos 1-3**; n=33, 10, 97, and 10 trials for DNp09, aDN2, MDN, and control flies, respectively). The same flies are shown in **c** and **d**. **(c)** Single neuron / region-of-interest (ROI) response magnitude to command-like DN stimulation. Each circle is scaled and color-coded to represent the maximum change in fluorescence (normalized Δ*F/F*) of one detected DN axon/ROI relative to the level of activity 1 s prior to stimulation. Small white dots are shown if the response magnitude is smaller than the 95% confidence interval of the mean across trials. The background image is a standard-deviation projection across time of raw fluorescence microscopy data. **(d)** Trial-averaged single neuron/ROI responses across time, aligned to stimulus onset and ordered by response magnitude. Data are color-coded according to the magnitude of activity, or white if the response is smaller than the 95% confidence interval of the mean. Indicated are the number of neurons/ROIs with a positive response magnitude larger than the 95% confidence interval of the mean across trials (horizontal red line). **(e-f)** A **(e)** registered overlay or **(f)** density visualization of the data from multiple flies analyzed in the same manner as in **c**. The number of flies and trials are the same as in **a**. **(g)** Statistical comparison of the number of activated neurons/ROIs (i.e., the red dashed line as in **d**) using Mann-Whitney-U tests (p-values: DNp09 vs. control = 0.018, aDN2 vs. control = 0.040, MDN vs. control = 0.008). **(h)** Statistical comparison of the fraction of activated neurons (i.e., data from **g** divided by the total number of visible neurons/ROIs per fly) using Mann-Whitney-U tests (p-values: DNp09 vs. control = 0.018; aDN2 vs. control = 0.040; MDN vs. control = 0.008). **(i)** Statistical comparison of the strength of activation (i.e., the sum of normalized Δ*F/F* for positively activated neurons) using Mann-Whitney-U tests (p-values: DNp09 vs. control = 0.018; aDN2 vs. control = 0.040; MDN vs. control = 0.008). Error bars in (g)-(i) represent 95% confidence interval of the mean.

When we trial-averaged the change in neural activity upon optogenetic stimulation of command-like DNs, we observed a clear increase in GNG-DN activity during the stimulation of any of the three sets of command-like DNs in individual animals: DNp09 **(Supplementary Video 1)**, aDN2 **(Supplementary Video 2)**, and MDN **(Supplementary Video 3) (Fig. 2b-d)**. This was also consistent across multiple animals **(Fig. 2e-f)**. Importantly, we did not observe such a pronounced recruitment of GNG-DNs in control animals lacking a spGAL4 transgene **(Supplementary Video 4, Fig. 2b-f, far-right)**. This confirms that these populations became active due to command-like DN stimulation. Statistical comparisons confirmed that, for all three sets of command-like DNs, the number and fraction of GNG-DNs activated were significantly higher than for control animals **(Fig. 2g-h**; DNp09: p=0.018, aDN2: p=0.040, MDN: p=0.008).

Interestingly, we found that GNG-DNs were recruited in a spatially distinct manner across the cervical connective depending on which command-like DNs were optogenetically activated **(Fig. 2e-f)**. The activation of forward walking (DNp09) and anterior grooming (aDN2) command-like DNs increased activity among DNs localized to distinct regions of the medial cervical connective: the entire dorsal-ventral axis for forward walking, and the central and ventral connective for grooming. Activation of backward-walking command-like DNs led to weaker GNG-DN recruitment in the central connective.

We quantified the strength of GNG-DN recruitment as the summed responses of neurons that were positively activated during optogenetic stimulation **(Fig. 2i)**. The summed response was significantly higher for the stimulation of command-like DNs compared with control animals (DNp09: p=0.018, aDN2: p=0.040, MDN: p=0.008). Intriguingly, we also observed a recruitment gradient among command-like DNs: DNp09 stimulation resulted in very strong recruitment of GNG-DNs, aDN2 slightly weaker recruitment, and MDN the weakest. We note that reduced GNG-DN activity **(Fig. 2b-f, blue neurons and regions)** likely does not reflect inhibition (which is not robustly observed using calcium imaging) but rather reduced drive in DNs that were previously active during spontaneous forward walking prior to optogenetic stimulation. Consistent with this, additional experiments in animals that were resting prior to optogenetic stimulation showed fewer DNs with reduced activity **(Supporting Information File 1)**.

To address the possibility that recruitment of GNG-DNs by optogenetic stimulation may be non-ethological rather than reflecting a natural process, we compared the activity of GNG-DN populations in the same animal both during optogenetic stimulation and during the corresponding natural behavior. Specifically, in individual animals we compared neural activity during DNp09 stimulation to bouts of spontaneous forward walking **(Extended Data Fig. 2a, Supplementary Video 5)**, aDN2 stimulation to air puff-induced anterior grooming **(Extended Data Fig. 2b, Supplementary Video 6)**, and MDN stimulation to spontaneous backward walking on a cylindrical treadmill **(Extended Data Fig. 2c, Supplementary Video 7, see Methods)**. In each of these cases, we observed that populations of GNG-DNs were recruited during both optogenetic and natural conditions. For backward walking, these patterns were largely similar across optogenetic and natural conditions **(Extended Data Fig. 2c)**. However, for forward walking **(Extended Data Fig. 2a)** and to a lesser extent for anterior grooming **(Extended Data Fig. 2b)**, there were some differences. DNp09 stimulation consistently and strongly activated a small subset of DNs located in the medial-dorsal and medial-ventral connective. These were not active during spontaneous forward walking **(Extended Data Fig. 2d-f)**. However, the remaining largest fraction of DNs were active in a similar fashion during optogenetic stimulation and during spontaneous walking **(Extended Data Fig. 2e, white region)**.

Taken together, these data reveal that the stimulation of command-like DNs leads to the recruitment of many additional DNs in the GNG in a manner that is similar to DN population activity during natural behaviors. This framework can reconcile the two observations that only a few command-like neurons are sufficient to drive behaviors but that larger DN populations are active during spontaneous behaviors.

### Command-like DNs connect to recurrent DN networks in the brain

The functional recruitment of GNG-DNs by command-like DNs could arise from a variety of circuit mechanisms. Broadly speaking, it might either result from direct, monosynaptic excitatory connections, or indirect polysynaptic connectivity via local brain circuits, or via ascending neurons in the VNC. To investigate these possibilities, we examined DN connectivity within the female adult fly brain connectome ^3,42^. There, we found our three sets of command-like DNs—DNp09, aDN2, and MDN **(Fig. 3a)**—and identified all of their downstream DN partners. We observed that each command-like DN has direct, monosynaptic connections to DNs with somata in both the cerebral ganglia (CG, purple) and the GNG (orange) **(Fig. 3b)**.

**Fig. 3:**
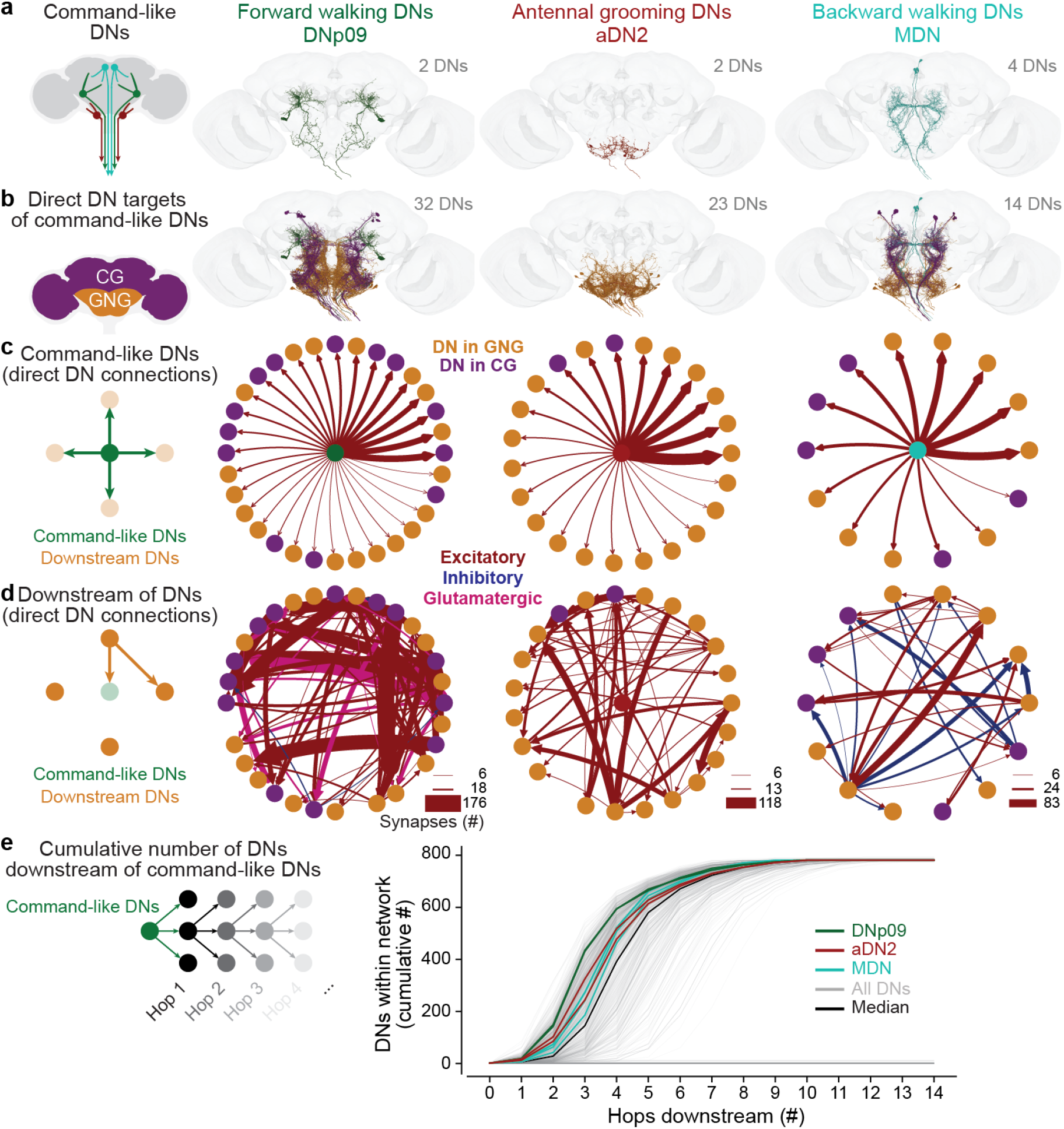
Command-like DNs synapse onto other DNs, forming larger DN networks. **(a)** The neuronal morphologies of three sets of command-like DNs—**(left)** DNp09, **(middle)** aDN2, and **(right)** MDN in the female adult fly brain connectome ^3^. **(b)** The location and morphologies of DNs directly (monosynaptically) targeted by command-like DNs. DNs are color-coded based on their cell body localization in the GNG (orange) or CG (purple). Command-like neurons are color-coded as in **a**. **(c)** Command-like DNs form monosynaptic excitatory connections to downstream DN targets. Edge weights reflect the number of synapses as shown in **d**, with consistent scaling across all plots. Edge colors denote excitatory (red), inhibitory (blue), or glutamatergic (pink) which can be excitatory or inhibitory depending on receptor type ^75^. DNs are color-coded as in **b**. **(d)** Network connectivity among downstream DNs shows strong recurrence and minimal feedback to command-like DNs (only in aDN2). **(e)** The cumulative number of downstream DNs that three sets of command-like neurons— DNp09 (green lines, 2 DNs), aDN2 (red lines, 2 DNs), MDN (cyan lines, 4 DNs) connect to across an increasing number of DN-DN synapses (‘hops’). This is compared to the number of DNs accessible over an increasing number of hops for all DNs (grey lines) and the median of all DNs (black line). The maximum number of recruited neurons (*∼* 800) is smaller than the total number of DNs because 455 neurons receive inputs from maximally one other DN. Note that many DNs do not connect to any other DN, even across 14 hops.

Based on predictions from EM images, our command-like DNs are cholinergic ^3,43^ and the connections they form with downstream DNs are most likely excitatory **(Fig. 3c, red arrows)**. These connections are predominantly feedforward with only sparse feedback connections for aDN2 **(Fig. 3d)**. By contrast, among downstream DNs, we observed strong recurrent interconnectivity, including some inhibition **(Fig. 3d, blue arrows)**. Notably, command-like DNs connect to a variable numbers of downstream DNs that mirrors the differential recruitment of GNG-DNs in our functional imaging experiments **(Fig. 2i)**: those for forward walking (DNp09) have the most downstream DNs (32), while those for antennal grooming (aDN2) have fewer (23), and those for backward walking (MDN) have the fewest (14). This ordering also holds for multi-synaptic connections to downstream DNs **(Fig. 3e)**. All three command-like DNs, in particular DNp09, form more short connections (one to six synapses away) to the remaining DN population than the median DN **(Fig. 3e)**. These data support a mechanism whereby command-like DNs engage additional DN populations in the brain via direct excitatory connections.

### Behavioral requirement for DN population recruitment

We next asked to what extent the recruitment of additional DN populations is necessary for command-like DNs to drive complete behaviors. We can envision at least four possibilities. First, it may be that these additional DNs are required because command-like DNs only control a small subset of the required behavioral kinematics. Second, additional DNs may be modulatory, controlling behavioral vigor or persistence. Third, additional DNs may ‘gate’ a behavior that is initiated by command-like DNs. Fourth, DN recruitment may be relatively inconsequential: activating command-like DNs alone may be sufficient to drive a complete behavior.

To distinguish between these possibilities, we needed to stimulate command-like DNs while preventing the recruitment of additional DN populations. We achieved this by carefully decapitating flies and sealing their exposed neck. It has been shown that flies can survive and generate behaviors for hours following decapitation ^44^. In this way we could identify which elements of behavioral kinematics directly result from optogenetic activation of command-like DNs alone (i.e., a low-dimensional signal coming from the brain), without the recruitment of other DNs in the brain **(Fig. 4a, right)**. Notably, a less invasive approach—acute optogenetic inhibition of GNG-DNs using the anion channelrhodopsin, GtACR1^45^ (Dfd-LexA *>* LexAop-GtACR1)—would achieve inhibition of only a fraction of all DNs and, when tested, caused animals to groom at even low light intensities, obstructing any analysis of command-like DN-driven behaviors.

**Fig. 4:**
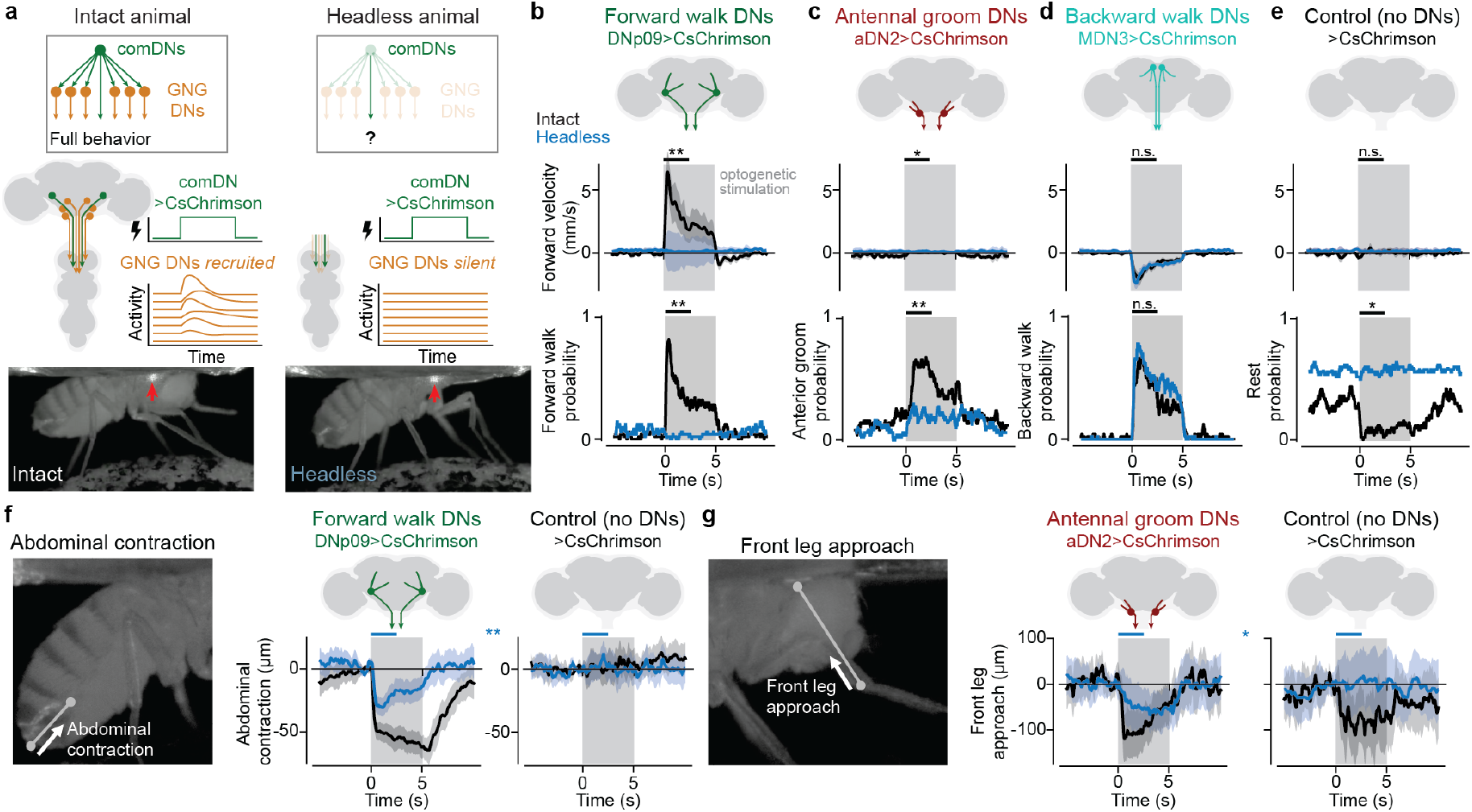
Recruited DN networks are required for forward walking and grooming, but not for backward walking. **(a)** In intact animals, the activation of a command-like DN (green) recruits other DNs (orange) and leads to the execution of a complete behavior (left). In headless flies, the axons of command-like DNs (green) can still be activated in the VNC. However, other DNs (orange) cannot be recruited in the brain and will remain silent (right). This comparison between intact and headless animals allows one to isolate the contribution of full DN networks versus command-like DNs alone to behavioral output. **(b-e)** Forward walking velocities and behavior probabilities for **(b)** DNp09, **(c)** aDN2, **(d)** MDN, or **(e)** control flies. Mann-Whitney-U tests compare the difference between the means of the first 2.5 s of optogenetic stimulation across intact (black traces) versus headless animals (blue traces). **(f)** DNp09 stimulation in both intact and headless animals leads to abdominal contractions. This is quantified as the change in Euclidian distance between the anal plate and ventral side of the most posterior stripe compared to 1 s prior to stimulation. Mann-Whitney-U test compares the mean of the first 2.5 s of stimulation (blue bars) for headless DNp09 versus headless control animals (two blue traces). **(g)** aDN2 stimulation in both intact and headless animals leads to front leg approach. This is quantified as the change in Euclidian distance between the front leg tibia-tarsus joint and the neck compared to 1 s prior to stimulation. Mann-Whitney-U test compares the first 2.5 s of stimulation (blue bars) between headless aDN2 and headless control animals (two blue traces). All plots in **b**-**g** show data from 5 flies with 10 trials each. Statistical tests compare the trial mean across different flies. *** = p*<*0.001, ** = p*<*0.01, * = p*<*0.05, n.s. = p*>*0.05. For exact p-values, see Methods.

We compared the behaviors of intact and headless animals upon optogenetic activation of command-like DNs. As in our previous experiments, stimulation of DNp09, aDN2, and MDN in intact animals drove forward walking **(Supplementary Video 8)**, anterior grooming **(Supplementary Video 9)**, and backward walking **(Supplementary Video 10)**, respectively (**Fig. 4b-d, black traces)**. Control animals with no DN driver did not reliably generate specific behaviors upon laser stimulation but were more aroused **(Supplementary Video 11)** resulting in a decreased probability of resting **(Fig. 4e, black traces)**.

After decapitation of the same animals, we found that the activation of MDN in headless flies still drove backward walking. This confirms that decapitation does not trivially impair movement generation (**Fig. 4d**; *p* = 0.265 comparing the backward walking probabilities of headless versus intact flies). By contrast, decapitation had a different effect on the other two command-like DNs: DNp09 and aDN2 stimulation in headless animals did not generate forward walking (**Fig. 4b**; *p* = 0.006) or anterior grooming (**Fig. 4c**; *p* = 0.006), respectively. Importantly, headless animals could still behave. Outside of optogenetic stimulation periods we observed episodes of spontaneous grooming in headless flies that resembled those generated by intact animals. This confirms that local VNC circuits required to generate coordinated movements were still intact. Additionally, although optogenetic stimulation of DNp09 and aDN2 in headless flies did not drive complete forward walking or anterior grooming, respectively, it reliably elicited more subtle movements: stereotyped abdomen contraction for DNp09 (**Fig. 4f**; *p* = 0.006 comparing headless DNp09 versus headless control animals) and front leg approach for aDN2 animals (**Fig. 4g**; *p* = 0.030 comparing the distance between the tibia-tarsus joint and neck in headless aDN2 versus headless control animals).

In summary, these data lead us to posit that differences in optogenetically-driven behaviors between intact and headless flies result from the failure to recruit additional, downstream DN pathways in the brain. These results also show that the functional recruitment of populations of DNs is necessary for command-like DNs to drive more flexible (DNp09 and aDN2), but not stereotyped (MDN) behaviors, suggestive of different modes of DN-driven behavioral control that we next set out to explore.

### Descending neuron connectivity predicts the necessity for DN recruitment to drive behavior

Our results thus far reveal a correlation between three properties of command-like DNs **(Fig. 5a, top)**: the functional recruitment of other DNs **(Fig. 2)**, monosynaptic connectivity to downstream DNs **(Fig. 3)**, and the necessity of downstream DNs to generate complete optogenetically-driven behaviors **(Fig. 4)**. For example, for DNp09, optogenetic stimulation activates many other DNs, DNp09 has many downstream DN partners, and they cannot drive forward walking in headless animals. By contrast, for MDN, optogenetic stimulation activates fewer GNG-DNs, MDN has fewer direct downstream DN partners, and they alone are sufficient to drive backward walking in headless animals. Thus, we propose that command-like DNs lay on a continuum between ‘broadcaster’ DNs like DNp09, which have a large number of downstream DNs that are recruited and required to execute flexible behaviors, and ‘stand-alone’ DNs which do not have any downstream DNs and are by themselves sufficient to drive stereotyped movements **(Fig. 5a, gray box)**. This hypothesis suggests that the strength of a DN’s connectivity to other DNs is predictive of the behavioral outcome of optogenetic stimulation in intact versus headless animals. Specifically, broadcaster or stand-alone DNs should exhibit a strong or weak degradation of a flexible or stereotyped behavior, respectively, following decapitation **(Fig. 5a, light blue box)**.

**Fig. 5:**
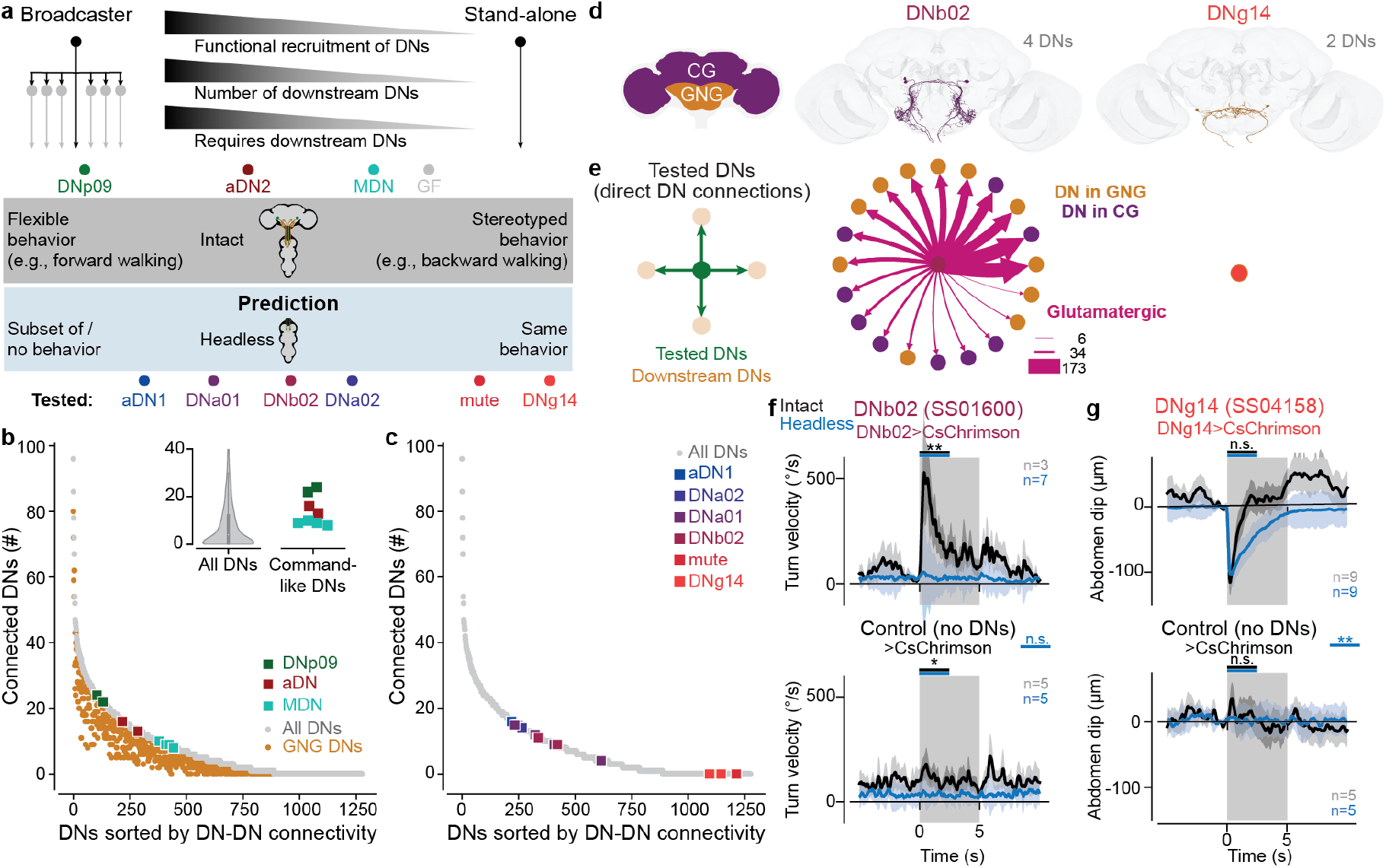
Network connectivity accurately predicts the necessity for other DNs and flexibility of DN-driven behaviors. **(a)** For the three command-like DNs investigated three properties covary: (i) the number of functionally recruited DNs, (ii) the number of directly connected downstream DNs, and (iii) the requirement of downstream DNs to generate a complete behavior. This implies a continuum across DNs that spans from ‘broadcaster’ DNs like DNp09—which recruit large networks of DNs that are required to drive flexible behaviors—to ‘stand-alone’ DNs—which recruit no other DNs to drive stereotyped behaviors. We include giant fiber neurons (GF, grey) in this category based on previous studies of headless animals ^46,47^ and their small number of downstream DNs. These findings predict the requirement and behavioral flexibility of other, untested DNs. Namely, that optogenetically activating a broadcaster DN with many directly connected downstream DNs should drive more flexible behaviors that are lost in headless animals. Conversely, optogenetically activating a stand-alone DN with few directly connected downstream DNs should drive relatively simple behaviors that are retained in headless animals. Schematized along this continuum are our three tested command-like DNs (DNp09, aDN2, and MDN), as well as six additional untested neurons for which we make connectome-based predictions: aDN1, DNa01, DNb02, DNa02, Mute, and DNg14. Their positions on the horizontal continuum are coarsely defined. **(b)** The number of all (grey circles) or GNG-based (orange circles) DNs monosynaptically downstream of each individual *Drosophila* DN. Command-like DNs of interest are color-coded as elsewhere. Inset shows the median, 25 %, and 75 % quantile of DNs targeted for all DNs (left violin plot) and the number of DNs targeted by three sets of command-like DNs (DNp09, aDN2, MDN). **(c)** The number of DNs directly downstream of six sets of additional DNs (color-coded circles as in **a**) for which we make connectome-based predictions. All DNs are shown for reference as in **b** (grey circles). **(d)** The morphology in the female adult fly brain connectome of two sets of DNs (DNb02, DNg14) for which we test connectome-based predictions. **(e)** Monosynaptic connectivity for the two tested DNs. DNb02 has many connections, akin to the ‘broadcaster’ DN class, whereas DNg14 does not connect to any other DNs in the brain, akin to the ‘stand-alone’ DN class. Edge weights denote the number of synapses. Edge colors denote excitatory (red), inhibitory (blue), or glutamatergic (pink) which can be excitatory or inhibitory depending on receptor type ^75^. **(f)** Absolute turn velocity for DNb02 (top) and control (bottom) animals upon optogenetic stimulation. This magnitude of the turn velocity is not directed (i.e., leftward or rightward). **(g)** Abdomen dipping for DNg14 (top) and control (bottom) animals upon optogenetic stimulation. Abdomen dipping is quantified as the change in vertical position of the anal plate. In **f** and **g**, shown are data for intact (black traces) and headless (blue traces) animals. The number of animals are indicated for each condition. Each animal was optogenetically stimulated 10 times. Thus, the traces show the average and 95 % confidence interval of the mean across *n ∗* 10 trials. Shown are Mann-Whitney U tests comparing the trial mean of intact and headless animals (black bars) or comparing headless experimental with headless control flies (blue bars, result shown between top and bottom plot). *** = p*<*0.001, ** = p*<*0.01, * = p*<*0.05, n.s. = p*>*0.05. For exact p-values, see Methods.

To test these predictions, we sought additional broadcaster and stand-alone DNs by examining direct DN-DN connectivity for all DNs in the brain connectome. There, we observed a continuum of DN-DN connectivity for DNs across the entire brain **(Fig. 5b, gray)** that was also present in GNG-based DNs specifically **(Fig. 5b, orange)**. A few DNs have dozens of DN partners while hundreds of DNs have no downstream DNs. Our three sets of command-like DNs are in the middle of this continuum with higher connectivity than most DNs (median number of connected DNs: 4; MDN: 9; aDN2: 15; DNp09: 23; **Fig. 5b, inset**). Notably, consistent with our hypothesis, giant fiber neurons (GF) have only a few DN partners (three and four for the left and right GFs, respectively; **Fig. 5a, gray**) and are known to drive stereotyped, ballistic escape in both intact and headless animals ^46,47^. We selected six sets of DNs along this continuum **(Fig. 5c, colored squares)** by identifying those with available genetic driver lines ^14,15^ and that fulfilled additional criteria (see Methods). We optogenetically stimulated these sets of DNs in animals that were either intact or headless. Data from these six experiments confirmed our predictions: DNs driving more flexible behaviors in intact animals and with many downstream DNs lost their behavior in headless animals while DNs without any downstream partners elicited stereotyped movements that were maintained in headless animals. Among broadcasters, this was most profound for DNb02. These DNs synapse upon 20 other DNs **(Fig. 5d-e)** and drive turning in intact animals. In headless animals, DNb02 stimulation does not elicit turning (**Fig. 5f**, *p* = 0.001 comparing intact and headless flies) but instead drives flexion of the front legs at stimulation onset **(Supplementary Video 12)**. This is noticeable as a small spike in forward velocity in headless animals **(Extended Data Fig. 3c)**. Similarly, turning was lost in DNa01 **(Extended Data Fig. 3b, Supplementary Video 13)** and DNa02 **(Extended Data Fig. 3d, Supplementary Video 14)**) animals. aDN1 animals retained only partial behaviors— anterior grooming became uncoordinated front-leg movements in headless animals **(Extended Data Fig. 3a, Supplementary Video 15)).**

Among stand-alone DNs, the maintenance of stereotyped behaviors was most clear for DNg14. These DNs do not directly synapse upon any other DN **(Fig. 5e)** and their activation triggers a subtle dip and vibration of the abdomen in both intact and headless animals (**Fig. 5g**, *p* = 0.144, **Supplementary Video 16**). Similarly, ovipositor extension was retained in headless Mute animals **(Extended Data Fig. 3f, Supplementary Video 17)**).

Thus, our hypothesis that DN downstream connectivity is predictive of the kind of DN-driven behavior (flexible versus stereotyped) and its requirement for network recruitment is supported by experiments on ten DNs spanning the continuum from those driving flexible behaviors (DNp09-induced forward walking, DNa01/DNa02/DNb02 induced turning, and aDN1/aDN2-induced grooming) to those driving stereotyped (giant fiber-induced escape jump ^47^, and MDN-induced backward walking) behaviors and movements (DNg14-induced abdominal dipping/vibration, and Mute-induced ovipositor extension).

### DN networks cluster as a function of associated behaviors

Our investigation of the brain connectome revealed that DN-DN connectivity lies on a continuum: a few DNs have very high, hub-like connectivity (e.g., *>*80 downstream DNs), while 567 (44 %) target only two or fewer DNs **(Fig. 5b)**. Hub-like DNs of high-degree connectivity suggest an overall structure of DN networks that would have implications on how information flows between neurons. Therefore, we next examined the large-scale structure of the entire DN network. To do this, we compared the DN network derived from the fly brain connectome with a shuffled DN network having the same number of neurons and the same number of overall connections between neurons, but with individual connections randomly assigned. We found that the connectivity degree distribution (i.e., the distribution of how many other DNs each DN connects to) is indeed dramatically different (*R*^2^ = *−*0.04 comparing connectivity distributions) between the original DN network (**Fig. 6a, black**) and the shuffled network (**Fig. 6a, red**). This is largely because very strongly connected DNs (*>*30 partners) and very weakly connected ones (*<*5 partners) only appear in the original connectome DN network but not in the shuffled network. The original DN network can be fit better by an exponential (*R*^2^ = 0.92, green) or a power law (*R*^2^ = 0.79, blue) degree distribution, indicative of intrinsic structure. A power law connectivity degree distribution is the defining feature of a scale-free network ^48,49^ and hints at an underlying structure of DNs linked through well-connected ‘hub’ neurons.

**Fig. 6:**
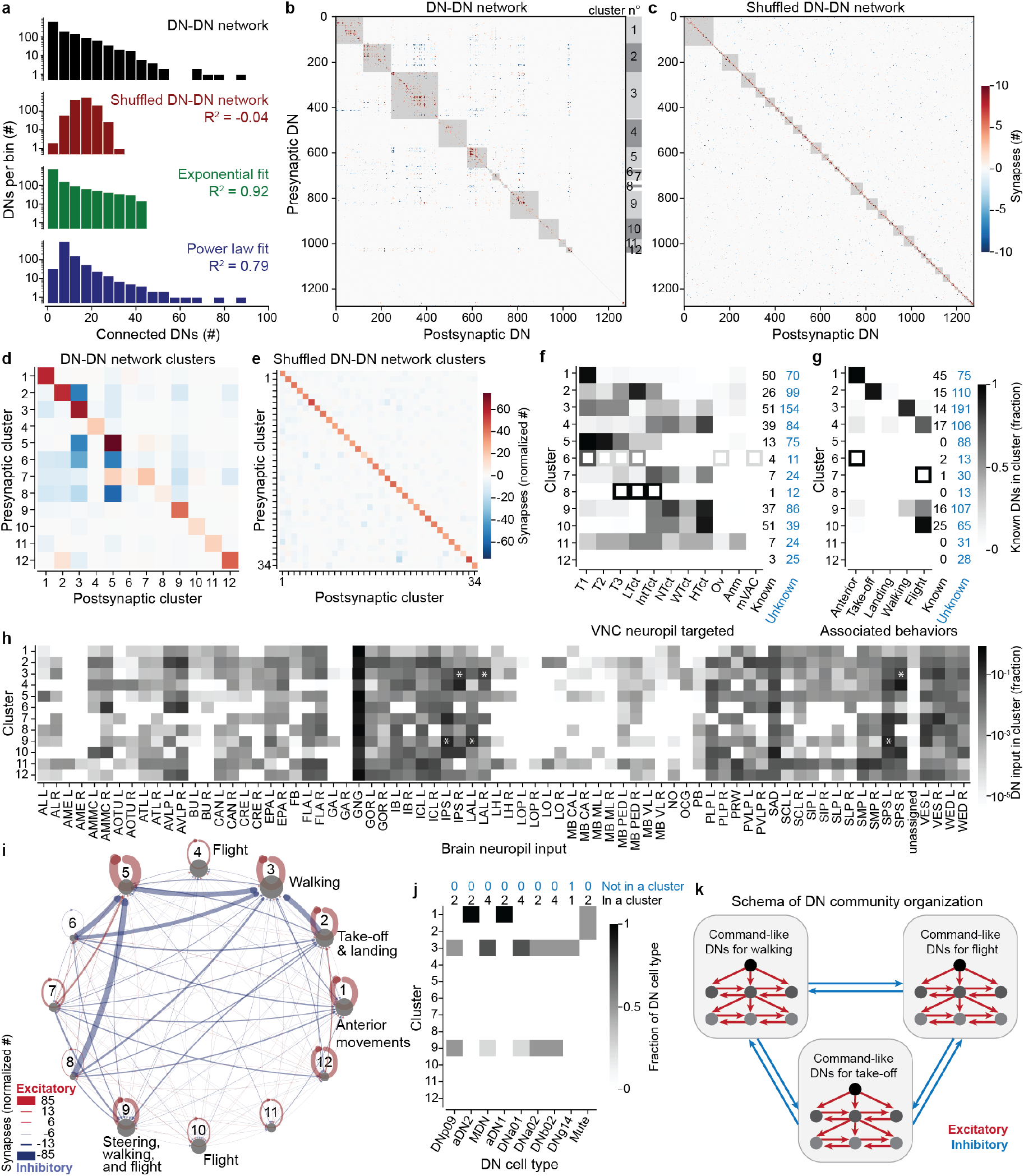
Networks of DNs for similar behaviors excite one another and inhibit those for other behaviors. **(a)** The connectivity degree distribution of (black) the DN-DN network as a histogram (bin width 5) on a logarithmic scale and (red) the same data after shuffling individual connections between DNs. Shown as well are the best exponential (green) or power-law degree distribution (blue) fits to the DN-DN network data. **(b)** A community detection algorithm applied to the DN connectivity matrix yields large clusters (grey squares). DNs are sorted based on detected clusters. Excitatory (red) and inhibitory (blue) connectivity between individual DNs from row (presynaptic) to column (postsynaptic) are color-coded. Numbers on the right side map onto cluster numbers in **d,f,g,h,i,j**. **(c)** Equivalent analysis to **b**, but for a network with shuffled DN-DN network connectivity. **(d)** The number of synapses between any two clusters normalized by the number of DNs in the postsynaptic cluster. Only clusters with 10 or more DNs are considered. Synapse weights are summed such that one excitatory synapse and one inhibitory synapse is shown as no synapse. **(e)** Equivalent analysis to **d**, but for the shuffled network in **c**. **(f)** For each cluster, the fraction of known DNs projecting to different neuropil regions within the VNC (T1,T2,T3: leg neuropils, LTct: lower tectulum, IntTct: intermediate tectulum, NTct: neck tectulum, WTct: wing tectulum, HTct: haltere tectulum, Ov: ovoid, Anm: abdominal neuromere, mVAC: medial ventral association center) is shown. Data are taken from ^9^ (**Supporting Information File 1**). The number of known (black) versus unknown (blue) DNs per cluster is indicated (right). Empty squares indicate clusters containing fewer than five known DNs. **(g)** For each cluster, the fraction of known DNs associated with different behaviors (anterior movements, take-off, landing, walking, flight) is shown. The number of known (black) and (blue) unknown DNs per cluster is indicated (right). Data are taken from the literature (**Supporting Information File 2**). Empty squares indicate clusters containing fewer than five known DNs. **(h)** For each cluster, the fraction of DN input synapses from different brain neuropils. Asterisks indicate a left-right imbalance among inputs to clusters 3 and 9 from the IPS, SPS and LAL. Note that these data are shown on a log scale. Neuropil names are described in Methods. **(i)** A network visualization of clusters in **d**. Overlaid are associated behavior annotations from **g**. Edge weights indicate the relative strength of excitatory (red) or inhibitory (blue) connectivity. **(j)** The distribution of optogenetically investigated DNs across DN clusters. The number of DNs inside (black) and outside (blue) the 12 clusters is indicated (top). **(k)** A summary model of the proposed organization among DN networks. There are predominantly excitatory (red) connections within each DN cluster. Each cluster may have its own hierarchical network sub-structure, with command-like DNs recruiting other DNs in a excitatory fashion. There are inhibitory (blue) connections between DN clusters.

Inherent structure within this network may be indicative of sub-networks, or clusters, with unique properties. In the case of DNs, our initial analysis of command-like DNs downstream partners suggest the existence of non-overlapping networks for forward walking (DNp09) versus anterior grooming (aDN2). To further explore this possibility, we identified clusters of DNs within the entire fly brain and applied the Louvain method—a community detection algorithm ^50^—to the undirected network of DNs (i.e., connections between two neurons are scaled by their synaptic strength and neurotransmitter identity, but the directionality of the connection is not taken into account; see Methods). Indeed, we could reliably identify multiple clusters of DNs with strong inter-connectivity **(Fig. 6b, gray boxes)**. Importantly, when we applied the same algorithm to our shuffled network with the same size, same number of neurons, same connection weights, but randomly shuffled connections, we only inconsistently found small clusters **(Fig. 6c, gray boxes)**. This was clear by quantifying the number of DNs in the five largest clusters for the original DN-DN network (726*±*42 neurons) versus the shuffled DN-DN network (581 *±* 51 neurons; mean *±* std, *p <* 0.001 comparing 100 repetitions of the Louvain method).

Within clusters we observed predominantly strong excitatory connections **(Fig. 6d, diagonal elements)**. By contrast, between clusters we observed predominantly inhibition **(Fig. 6d, off-diagonal elements)**, localized between specific clusters. In the shuffled DN-DN network, this inhibition was weaker and more uniformly distributed **(Fig. 6e, off-diagonal elements)**.

Distinct excitatory clusters could imply parallel pathways of descending neurons with distinct anatomical and/or functional properties. We examined this possibility by first investigating whether DN clusters (with similar connectivity in the brain) connect to similar targets in the VNC. Specifically, we studied the projections of known DNs ^14,20^ within the VNC connectome of an adult male fly ^9^. This analysis revealed very specific connectivity patterns including, for example, that cluster 1 projects predominantly to regions controlling the front leg (T1 neuropil), cluster 2 predominantly to the lower tectulum (LTct), clusters 3 and 5 most strongly to the leg neuropils (T1, T2, T3) and clusters 4, 7, 9 and 10 predominantly to dorsal neuropil regions involved in wing, haltere, and neck control (WTct, HTct, NTct) **(Fig. 6f)**.

These results strongly imply that excitatory DN clusters may also regulate distinct behaviors. We examined this possibility by identifying 132 known DNs which are shown or predicted to be involved in anterior movements, walking, take-off, flight, and landing (see **Supplementary Data File 2** and Methods for details). Indeed, we found that the clusters included DNs with known links to specific behaviors **(Fig. 6g)**. For example, as might be expected, DNs related to anterior grooming were predominantly in cluster 1 which targets T1 (e.g., DNg10^19^, DNg12^19^, aDN1^18^, aDN2^18^). All DNs involved in take-off and, more weakly, landing were in the cluster 2 (e.g., giant fiber/DNp01^10^, DNp02^30^, DNp04^30^, DNp10 for landing ^13^, DNp11^30^, DNp35^9^). Many DNs related to walking were in cluster 3 with strong projections to T1-T3 (e.g., DNp09^17^, DNa01^51^, DNa02^52^, DNb02^53^, DNg13^9^, MDN ^20^). This cluster also receives strong inhibition, particularly from cluster 2 related to take-off **(Fig. 6i)**. Cluster 5 also mainly projects to the leg neuropils (T1-T3), but the behavioral phenotype of its constituent DNs are unknown. DNs related to flight were also localized to clusters 4 (e.g. DNa10^9^, DNg02^29^, DNp20^54^) and 10 (e.g. DNg02^29^, DNg07^29^, DNp31^9^). Some additional flight-related neurons involved in steering are in cluster 9 (e.g. DNp03^9^, DNg02^29^), grouped with walking DNs known to be involved in turning.

The command-like DNs we studied experimentally were part of behavioral consistent clusters as well **(Fig. 6j)**. aDN1, aDN2 in the ‘anterior movement’ cluster 1 and DNp09, MDN, DNa01, DNa02, DNb02 in the ‘walking’ or ‘steering’ clusters 3 & 9, with neurons in the right hemisphere being assigned mainly to cluster 3 and those of the left hemisphere being assigned to cluster 9. This split of DNs associated with walking between clusters 3 and 9 was due to differences in connectivity between the two brain hemispheres, both in terms of bilateral symmetry in the brain as well as from the localization of the inputs coming from the inferior posterior slope (IPS), superior posterior slope (SPS) and the lateral accessory lobe (LAL) **(Fig. 6h, white asterisks)**.

Taken together, these data support a model in which complete behaviors are orchestrated by specific excitatory DN subnetworks which, in turn, inhibit other subnetworks driving potentially conflicting behaviors **(Fig. 6k)**.

## Discussion

Here, by combining optogenetic activation, functional imaging, and brain connectome analysis, we have resolved two seemingly conflicting observations: The activation of a few command-like DNs is sufficient to drive complete behaviors like forward walking, nevertheless many DNs are active when the same behavior is generated spontaneously. To explain this discrepancy, we propose a model in which command-like DNs recruit additional DN networks. This network recruitment is required for flexible forward walking but less critical for more stereotyped backward walking. Indeed, brain connectivity analyses suggest that all DNs generally lie along a continuum in which those driving more flexible behaviors recruit and require the activity of large DN networks, whereas those driving stereotyped behaviors and movements have dispensable connections to only a few downstream DNs. This model makes predictions which we confirmed through experiments on additional DNs. Finally, DN networks at a larger scale form excitatory clusters with specific behavioral roles and downstream targets in the motor system. These functionally distinct clusters inhibit one another, providing a potential mechanism for action selection in the brain.

### Circuit mechanisms giving rise to DN networks

There are a number of circuit motifs that could give rise to DN-DN interactions. Although we focus on monosynaptic connectivity, we have also shown that command-like DNs (DNp09, aDN2 and MDN) ultimately reach—and may potentially activate—hundreds of other DNs within only a few synapses. Thus, DN populations recruited by the activation of command-like DNs in our functional studies may also be multiple synapses away. Although the temporal limitations of calcium imaging preclude the ability to distinguish between mono- or polysynaptic recruitment, in the future it may be possible to map the identity of recruited DNs onto the connectome by taking advantage of volumetric anatomical recordings of the cervical connective and anterior VNC along with computational approaches to register functional recording data to an anatomical template ^55^.

Most of the monosynaptic connections between DNs are in the GNG. Akin to the mammalian brain stem, the GNG is a multi-sensory processing region ^15,56,57,58^ that controls head and proboscis movements ^3,59^. The presence of numerous DN inputs and outputs also make the GNG well-positioned as a locus for action selection ^60^. In line with this possibility, we found predominantly excitatory clusters of DNs that mediate distinct behaviors and inhibit one another.

Interestingly, the GNG also receives a large number of inputs from ascending neurons (ANs) that project from the VNC to the brain ^61,62^. Among these are a set of ANs involved in deciding between locomotion and feeding ^63^. Connections from ANs may thus allow DNs to integrate motor information from the VNC potentially to regulate switching during a sequence of actions. DN-DN recruitment could also potentially arise indirectly through ANs in a DN-AN-DN ‘zigzag’ motif that has previously been observed in low numbers in the *Drosophila* larval connectome ^64^ (i.e., a DN targets an AN in the VNC which projects back to the brain and to a different DN). Our headless fly experiments cannot discriminate the contribution of ANs (decapitation eliminates both DN-DN and putative DN-AN-DN connections). ANs encompass 17 % of all DN post-synaptic partners in the VNC ^9^ and 10 % of all DN pre-synaptic partners in the brain. Efforts aiming to bridge the existing brain and VNC connectomes ^3,8,65^ and to generate complete adult nervous system connectomes will further reveal the extent of DN-AN-DN circuit motifs in adult flies.

Alternatively, DN recruitment might arise indirectly via sensory feedback during a change in behavioral state: Active DNs may drive a new behavior which results in sensory feedback that in turn may be transmitted through ANs to influence other DNs. We would expect to see such sensory-induced DN activation in spontaneous occurrences of the behavior. However, we instead observed that, in general, fewer DNs are strongly activated during spontaneous behaviors compared with during optogenetically elicited behaviors. This argues that strong GNG-DN activation is a specific response to optogenetic stimulation of command-like DNs. In particular, we observed that the 10 most active neurons during DNp09 stimulation are not active during spontaneous forward walking. These results suggest that DN recruitment likely does not result from sensory feedback arising during optogenetically-induced changes in behavioral state.

In addition to excitatory interactions among DNs, on a larger scale, we observed that DN clusters related to distinct behaviors inhibit one another. Direct DN-DN connectivity is rare in the VNC (only 2 % of all DN post-synaptic partners in the VNC are DNs ^9^) compared to strong and prominent DN-DN connections within the brain (32 % of all DN post-synaptic partners in the brain are DNs). This inhibition is thus likely restricted to the brain because these DN clusters mostly project to different downstream targets within the VNC. Most DNs project either to the dorsal neuropils (wing, neck, haltere), or to the ventral leg neuropils. Only a few DNs, related to behaviors like take-off and landing—which require the coordination of both legs and wings—project to both of these regions ^9,14^. Additionally, a preliminary examination of how DNs in this study connect to one another in the male VNC connectome ^9,65^ shows that DNp09, giant fiber, DNa01, DNa02, and DNg14 do not synapse onto other DNs in the VNC. MDN and DNb02 connect to one and two DN cell types in the VNC, respectively. Thus, we can conclude that DN-DN interactions in the VNC are rare.

### The utility of DN networks

We have shown that precise stimulation of multiple classes of command-like DNs recruits activity in many additional DNs. Thus, the ‘command’ signal is not only conveyed directly to the VNC, but can also be sent to other brain neurons that, in parallel, convey additional descending signals. Interestingly, the strength of this DN recruitment varies across command-like DNs in a manner that correlates with behavioral flexibility as well as the necessity of these connections to generate a complete behavior. These results are most consistent with a descending control model in which most DNs drive relatively simple body part kinematics. Then, some privileged DNs (e.g., command-like DNs) directly recruit other DNs that, together, act as building blocks to construct a full behavior. This is in line with earlier observations in other insects that suggest that descending fibers may ‘act in consensus’ to assemble a complete behavior from different ‘subroutines’ ^66^. For example, DNp09 is connected to and requires the actions of a large number of DNs to drive elaborate movements of the six legs for goal-directed walking during courtship ^17^. Both DNp09 (for forward walking) and MDN (for backward walking) synapse upon DNa01 and DNa02, two other DNs involved in turning ^51,52^. Turning may therefore represent a behavioral building block that is used to construct both forward or backward walking. Notably, we found that DNa01 and DNa02 also synapse onto other DNs and that their activation alone (in headless animals) is not sufficient to drive turning. Thus, they may also sit higher in a hierarchy of DN recruitment activating yet other DNs which control specific movements required for turning. In the extreme case, such building blocks may be single degree-of-freedom movements like the lowering of the abdomen driven by DNg14—a pair of DNs that do not have any downstream DNs, but receive inputs from twelve upstream DNs.

These behavioral building blocks are analogous to ‘motor primitives’—fundamental elements for generating flexible behaviors in both vertebrates and invertebrates ^67^. In mammals it has been posited that complex behaviors are learnt by flexibly combining such motor primitives ^68^ potentially through a motor cortical network gain modulation mechanism ^69^ and in vertebrates, frogs can combine motor primitives stored in the spinal cord ^70^. Combining different motor primitives at the level of DNs may thus be a mechanism to ensure flexible descending control ^71^.

By contrast, other DNs can act alone to drive relevant downstream motor circuits and behaviors. These include the giant fiber which connects to a very small number of DNs and triggers a stereotyped and ballistic escape behavior. Similarly, MDNs with few downstream DNs drive backward walking. Neural activity during spontaneous and optogenetically-elicited backward walking was very similar, confirming that the control of backward walking is highly stereotyped. Both giant fiber and MDN are sufficient to drive their respective behaviors and do not rely on other DNs. Such a distinction between the mechanisms of descending control for flexible (walking) versus stereotyped (stridulation) behaviors has been proposed from studies in Orthoptera ^72^. This continuum of well-connected (DNp09) to largely disconnected (giant fiber) command-like DNs demonstrates that the classification of a DN as ‘command-like’ lies not only in its DN-DN connectivity. Instead, command-like DNs can drive complete behaviors through two distinct mechanisms: either by relying strongly on co-recruitment of other DNs in the brain or solely through their actions on VNC circuits.

In general, a descending control framework in which DNs can potentially recruit additional DNs, each of which control simple movements, provides an effective substrate for the evolutionary modification of behaviors (e.g., diversification of species-specific courtship displays), or the generation of entirely new behaviors through the *de novo* coupling or uncoupling of DNs. Thus, it is likely that a similar mechanism is leveraged for descending control in other species including mammals ^26,68^ and may be used to inspire the design of better artificial controllers in engineering and robotics ^73^.

## Methods

### Fly stocks and husbandry

All experiments were performed on female adult *Drosophila melanogaster* raised at 25°C and 50 % humidity on a 12 hr light–dark cycle. The day before optogenetic experiments (22-26 h before), we transferred experimental and control ^76^ flies to a vial containing food covered with 20*µl* of all transretinal (ATR) solution (100mM ATR in 100 %ethanol; Sigma Aldrich R2500, Merck, Germany) and wrapped in aluminium foil.

### Functional imaging and behavior experiments

We generated transgenic flies expressing LexAop-opGCaMP6s (generous gift from Orkun Akin ^77^) under the control of a Dfd-LexA driver (generous gift from Julie Simpson ^41^) and having a copy of UAS-CsChrimson (Bloomington ID 55135): **(Table 1, ID 1)** We also generated flies that additionally expressed LexAop-tdTomato (Bloomington ID 77139): **(Table 1, ID 2)**. For most experiments we used the flies lacking tdTomato expression.

**Table 1:** Transgenic fly lines generated in this study.

| ID | Chromosome X | Chromosome II | Chromosome III |
| --- | --- | --- | --- |
| 1 |  | 20xUAS-CsChr.mVenus (attP40),<br>13xLexAop-opGCaMP6s<br>(su(Hw)attP5) | DfdLexA / TM6B |
| 2 |  | 20xUAS-CsChr.mVenus (attP40),<br>13xLexAop-opGCaMP6s<br>(su(Hw)attP5) | 13xLexAop-CD4-tdTomato<br>(VK00033),<br>DfdLexA / TM6B |
| 3 |  | LexOp-myr-TdTomato / CyO | DfdLexA / TM6B |
| 4 |  | LexOP-H2B::mCherry / CyO | DfdLexA / TM6B |
| 5 | 20xUAS-CsChr.[mVenus]attP18 | VT023490-p65.AD | VT44845.GAL4DBD |
| 6 | 20xUAS-CsChr.[mVenus]attP18 | R76F12-AD | R18C11-DBD |
| 7 | 20xUAS-CsChr.[mVenus]attP18 | VT023490-p65.AD | R38F04-GAL4.DBD |

MDN-spGAL4 flies (also known as MDN3 from ^20^) were used to drive backward walking. aDN2-spGAL4 flies (also known as aDN2-spGAL4-2 from ^18^) were used to drive antennal grooming. DNp09 spGAL4 flies (from ^17^) were used to drive forward walking. Their genotypes are listed at the top of **Table 2**.

**Table 2:**
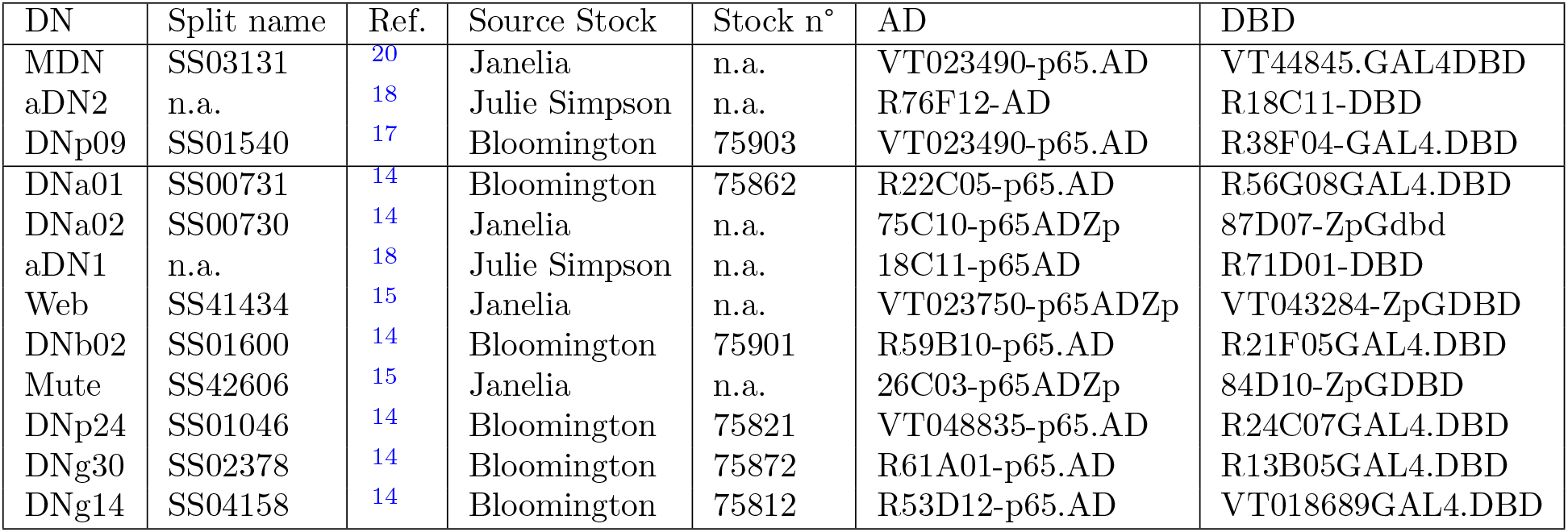
SpGAL4 lines used in this study.

For all experiments in **Fig. 2 and Fig. 4** we crossed those spGAL4 flies or wild type flies (PR - Phinney Ridge flies, Dickinson lab) with one of our stable transgenic driver lines for imaging **(Table 1, ID 1 or 2)**. For experiments in **Fig. 2** flies were 2-9 days post-eclosion at the time of experiments and experiments were performed at Zeitgeber time ZT 7-13. For experiments in **Fig. 4** flies were 2-9 days post-eclosion at the time of experiments and experiments were performed at Zeitgeber time ZT 4-7. For experiments in **Fig. 5, Extended Data Fig. 3** we crossed spGAL4 lines with 20XUAS-CsChrimson.mVenus (attP40) flies (Bloomington ID 55135). The exact genotypes of the split lines and the source stock are listed in the **Table 2**. All experiments were performed on flies 4-8 days post-eclosion (dpe), at Zeitgeber time ZT 4-7.

### Confocal imaging experiments

We generated flies with stable Dfd expression of membrane-targeted tdTomato, or nuclear-targeted mCherry based on flies generated by the Laboratory of Neural Genetics and Disease (McCabe laboratory, EPFL): **(Table 1, ID 3 and 4)**. For the three split-GAL4 driver lines targeting command-like DNs (MDN, DNp09, aDN2), we generated stable lines expressing UAS-CsChrimson: **(Table 1, ID 5,6, and 7)**. We crossed flies expressing a red fluorescent protein variant with flies expressing CsChrimson in a split GAL4 driver line to visualise the expression patterns using confocal imaging (**Extended Data Fig. 1**).

### Recording from DNs using a Dfd driver line

We leveraged a genetic-optical intersectional approach to selectively record from GNG-DNs. We chose to record from GNG-DNs because we found that 73 % of all DN-DN synapses in the brain connectome are in the GNG. Additionally, the GNG houses 60% of all DNs and 85% of all DNs have axonal output in the GNG ^14^. However the Hox gene Dfd does not include the entirety of all GNG-DNs as it excludes those driven by the Hox gene Sex combs reduced (Scr) ^78^. Sterne et al. ^15^ estimate that 550 cells in the GNG are Dfd positive and 1100 Scr positive, with only a small fraction expressing both. We show, for example, that aDN2, while localized to the GNG, is Dfd negative and thus most likely Scr positive **(Extended Data Fig. 1c)**. In our hands, functional imaging of DNs using an Scr driver line proved to be difficult because Scr expression extends into the neck and anterior VNC ^41^. Specifically, we observed strong expression of GCaMP in the tissues surrounding the thoracic cervical connective (potentially ensheathing glia ^79^), making it very hard to record the activity of DN axons. We expect that some Scr-positive DNs will be recruited by command-like DNs as for Dfd-positive DNs. Thus, we likely under-report the number of recruited GNG DNs.

### Limitations of selected spGAL4 driver lines

In addition to descending neurons, our aDN2 spGAL4 driver line (*aDN2-GAL4.2* ^18^) contains two groups of additional neurons. One pair is on the anterior surface of the brain and, based on our control experiments, likely is not or is only weakly activated by targeted optical stimulation of the neck (and not at all activated by the thoracic stimulation). Another is a set of neurons in the anterior VNC. Because other driver lines targeting aDN2 neurons with more, other off-target neurons have the same behavioral phenotype as our aDN2 driver ^18^, we are more confident that the effects we observe are due to stimulating the aDN2 neurons themselves.

Different studies report variable behavioral phenotypes for stimulating the DNp09-spGAL4 driver line: Some saw forward walking ^17^, while others observed stopping or freezing ^12,80^. We observed both: at our standard 10*µ*W optogenetic stimulation power, heterozygous animals mostly walked forward. Occasionally flies would only transiently walk forward and then stop, or alternate rhythmically between walking and stopping. With higher expression levels of CsChrimson (i.e., DNp09-spGAL4 *>* UAS-CsChrimson homozygous animals) we observed mostly freezing. We used heterozygous animals for our study.

### Immunofluorescence tissue staining and confocal imaging

We dissected brains and VNCs from 3–6 dpe female flies as described in ref. ^61^. For samples in Extended Data Fig. 1a,c, we fixed flies in 4 % PFA (Paraformaldehyde, 441244-1KG, Sigma Aldrich, Merck, Germany) in 0.1M PBS (Gibco PBS, pH 7.4, 10010-015, ThermoFisher Scientific, Switzerland). We then washed them six times for 10 min with 1 % Triton (Triton X-100, X100-100ML, Sigma Aldrich, Merck, Germany) in PBS (thereafter named 1 % PBST) at room temperature (RT). We then transferred them to a solution of 1 % PBST, 5 % Natural Goat Serum (Goat serum from controlled donor herd, G6767-100ML, Sigma Aldrich, Merck, Germany) and primary antibodies (see **Table 3**) and left them overnight at 4°C. We then washed the samples six times for 10 min with 1 %PBST at RT. We then transferred them to a solution of 1 % PBST, 5 % Natural Goat Serum and secondary antibodies (see **Table 3**) and left them for two hours at RT. We then washed the samples six times for 10 min with 1 % PBST at RT. We mounted the samples on glass slides using SlowFade (SlowFade Gold Antifade Mountant, S36936, ThermoFisher Scientific, Switzerland) and applied a cover slip. To space the slide and the cover slip, we applied a small square of two layers of double-sided tape at each edge. We sealed the edges of the coverslip with nail polish.

**Table 3:** List of antibodies used for immunofluorescence tissue staining.

| figure | type | AB name | dilution | company | AB ID |
| --- | --- | --- | --- | --- | --- |
| a,b,c | 1° | Anti-Bruchpilot (mouse) NC82 | 1:20 | Dev. Studies Hybridoma Bank | NC82 1ml supernatant |
| a,c | 1° | GFP Tag Rabbit | 1:500 | ThermoFisher | G10362 |
| a,b,c | 2° | Goat anti-Mouse Alexa 633 | 1:500 | ThermoFisher | A21052 |
| a,c | 2° | Goat anti-Rabbit Alexa 488 | 1:500 | ThermoFisher | A11008 |
| b | 1° | Living Colors DsRed | 1:500 | Takara | 632496 |
| b | 1° | Chicken to GFP Anti-GFP | 1:1000 | abcam | ab13970 |
| b | 2° | Goat Anti-Rabbit (Cy3) | 1:500 | abcam | ab6939 |
| b | 2° | Goat Anti-Chicken (Alexa 488) | 1:500 | abcam | ab150169 |

For samples in Extended Data Fig. 1b we fixed flies in 4 % PFA in PBS and transferred them to 1 % PBST and left them overnight at 4 °C. We then washed the samples 3 times for 15 min with 1 % PBST at RT. We then transferred them to a solution of 1 % PBST, 5 % Natural Goat Serum and primary antibodies (see **Table 3**) and left them overnight at 4 °C. We then washed the samples 3 times for 15 min with 1 % PBST at RT. We then transferred them to a solution of 1 % PBST, 5 % Natural Goat Serum and secondary antibodies (see **Table 3**) and left them overnight at 4 °C. We then washed the samples 3 times for 15 min with 1 % PBST at RT. We mounted the samples on glass slides using SlowFade and applied a cover slip. To space the slide and the cover slip, we applied a small square of two layers of double-sided tape at each edge. We sealed the edges of the coverslip with nail polish.

We imaged samples using a Leica SP8 Point Scanning Confocal Microscope with the following settings: 20x, 0.75 NA HC PL APO dry objective, 2× image averaging, 1024×1024 pixels, 0.52×0.52*µ*m pixel size, 0.57*µ*m z-step interval; green channel 488nm excitation, 500-540nm emission bandpass; red channel (imaged separately to avoid cross-contamination) 552nm excitation, 570-610nm emission bandpass; infrared channel (nc82, imaged in parallel to green channel) 638nm excitation, 650-700nm emission bandpass. We summed the confocal image stacks in the z-axis and rotated and translated the images to center the brain/VNC using Fiji ^81^.

### Optogenetic stimulation system and approach

We used a 640nm laser (Coherent OBIS 1185055 640nm LX 100mW, Edmund Optics, US) as an optogenetic excitation light source. We reduced the light intensity using neutral density filters (Thorlabs, Germany) and controlled the light intensity using mixed analogue and digital control signals coming from an Arduino with custom software. A digital signal was used to turn the laser on and off. An analogue signal (PWM output from Arduino and RC-low pass filtered) was used to modulate the power. Both of those signals were sent in parallel to the laser and an acquisition board and were recorded alongside the two-photon microscope signals using ThorSync 3.2 Software (Thorlabs, Germany). The light was directed towards the fly using multiple mirrors. Fine control of the target location was achieved using a kinematic mount (KM100, Thorlabs, Germany) and a galvanometric mirror (GVS011/M, Thorlabs, Germany). We manually optimized targeting of the laser onto the neck/thorax before each experiment. The light was focused onto the fly using a plano-convex lens with *f* = 75.0 mm (LA1608, Thorlabs, Germany) placed at the focal distance from the fly.

We note that, although command-like DNs have axon collaterals in the GNG, none of the command-like DNs in this study were among the DN populations we imaged: DNp09-spGAL4 and MDN-spGAL4 lines drive expression in neurons with cell bodies in the cerebral ganglia and not in the GNG **(Extended Data Fig. 1a)**. The DN cell bodies of the aDN2-spGAL4 line are within the GNG but do not overlap with Dfd driver line expression **(Extended Data Fig. 1c)**. Thus, we could be certain that any active DNs would be recruited through synaptic connections and not optogenetically. We identified laser light intensities that could elicit robust forward walking, anterior grooming, and backward walking **(Fig. 2a, Extended Data Fig. 1d)**.

We used different laser intensities to stimulate MDN (10 *µ*W), DNp09 (10 *µ*W), and aDN2 (20 *µ*W) animals, respectively, because 10 *µ*W stimulation power mostly causes aDN2 animals to stop **(Extended Data Fig. 1d)**. Activation of MDN in the head, neck, and thorax were sufficient to trigger backward walking **(Extended Data Fig. 1e)**. Although some tissue scattering of laser light can be expected, in control experiments we found that activation of the head capsule, but not the thorax, could strongly elicit forward walking in the brain’s “Bolt protocerebral neurons” (BPNs)— these neurons are known to drive robust and fast forward walking ^17^**(Extended Data Fig. 1f)**. 10 *µ*W stimulation was more specific than 20 *µ*W, which is why we selected 10 *µ*W stimulation for MDN and DNp09 as well as the spGAL4 lines tested in **(Fig. 5f-g, Extended Data Fig. 3)**. We regularly calibrated the laser intensity by measuring it with a power meter (PM100D, Thorlabs, Germany) and adjusting the analogue gain of the laser.

### *In vivo* two-photon calcium imaging experiments

We performed two-photon microscopy with a ThorLabs Bergamo II two-photon microscope augmented with a behavioral tracking system as described in ref. ^28^. Briefly, we recorded a coronal section of the thoracic cervical connective using galvo-resonance scanning at around 16Hz frame rate. Additionally, optogentic stimulation was performed as described above. We only recorded the green PMT channel (525 *±* 25 nm) because the red PMT channel would be saturated by red laser illumination of the fly. In parallel, we recorded animal behavior at 100 frames per second (fps) using two infrared (IR) cameras placed in front of and to the right of the fly.

Flies were dissected to obtain optical access to the VNC and thoracic cervical connective as described in ref. ^51^. Briefly, we mounted the fly to a custom stage by gluing its thorax and anterior head to the holder and removed its wings. Then, we opened the dorsal thorax using a syringe needle and waited for indirect flight muscles to degrade for *∼* 1.5 h. We pushed aside the trachea and resected the gut and salivary glands. For some flies, where the trachea was obstructing the view, we placed a V-shaped implant^82^ into the thoracic cavity to push the trachea to the side. We then placed the fly over an air-suspended spherical treadmill marked with a pattern visible on IR cameras for ball tracking (air flow at 0.6 l/min). While the fly was adapting to this new environment, the imaging region was identified and the optogenetic stimulation laser was centered onto the neck (*∼* 15 min).

We then recorded 10000 microscopy frames (around 10 min) while also recording behavioral data using cameras placed around the fly and presenting optogenetic stimuli. During a typical 10 min recording session, we presented 40 stimuli (5 s stimulation and 10 s inter-stimulus interval). Whenever the recording quality was still good enough (many neurons were visible and the fly still behaved healthily), we recorded multiple sessions to increase the number of stimulation trials.

### Investigating natural behaviors

In **Extended Data Fig. 2** we compared optogenetically-elicited neural activity to activity observed during natural behaviors: forward walking, anterior grooming, and backward walking. Natural forward walking is frequently spontaneously generated by the flies. By contrast, we needed to stimulate the antennae with 5 s puffs of humidified air to increase the probability of natural grooming **(Extended Data Fig. 2b)**. We provided humidified air puffs with an olfactometer (220A, Aurora Scientific, Canada) using the following parameters: 80 ml/min air flow, 100 % humidity, 5 s duration, 20 s inter-stimulus interval. In order to have humid air puffs (i.e., an abrupt change in flow rate) instead of a switch from dry air to humidified air—the default olfactometer configuration—we only connected the “Odor” tube to the final valve and not the “Air” tube. Furthermore, to increase the likelihood of spontaneous backward walking **(Extended Data Fig. 2c)**, we replaced the spherical treadmill with a custom cylindrical treadmill that we found increases the motivation to backward walk. Specifically, we designed a 10 mm diameter, 80 mg 3D printed wheel (RCP-30 resin) and printed it using stereolithography through digital light processing (Envisiontec Perfactory P4 Mini XL). This wheel was mounted on a low-friction jewel-bearing holder (ST-3D sapphire shafts, VS-40 sapphire bearings, Freudiger SA, Switzerland). We marked the sides of the wheel with IR-visible dots to facilitate IR camera tracking of the wheel and calculations of velocity to classify bouts of backward walking. When using the wheel, we used an additional third IR camera to the left of the wheel, where dot markers were visible.

### Behavioral experiments in headless animals

For behavioral experiments, we mounted flies to the same stages used for two-photon imaging, but without gluing the anterior part of the head to the holder. Then, without further dissection, we placed animals onto the spherical treadmill. After recording 10 trials of responses to optogenetic stimulation in intact animals, we decapitated the fly by inverting the holder and pushing a razor blade onto the neck. To achieve this, we mounted a splinter of the razor blade onto the tip of a pair of dissection forceps for finer control. We took care not to injure the fly’s legs and to make a clean cut without pulling out thoracic organs passing through the neck connective. To limit desiccation, we then sealed the stump of the neck with a drop of UV-curable glue. We only continued experiments on flies if their limbs were moving following decapitation. We then placed the headless flies onto the spherical treadmill and let them recover for at least 10 min. Then, we recorded 10 trials of responses to optogenetic stimulation on the spherical treadmill and 10 trials in which the fly was hanging from the holder without contacting the spherical treadmill. In experiments for testing connectome-based predictions, we slightly modified this experimental procedure. Because intact control animals become aroused by optogenetic stimulation, to avoid false positives and to discover behavioral phenotypes for less well-studied DNs, we attempted to reduce the spontaneous movements of flies. First, instead of 10 s between optogenetic stimulation trials, we used 20 s. Second, we filled the fly holder with room-temperature saline solution to buffer heating from IR illumination.

### Data exclusion

We manually scored the quality of neural recordings (signal-to-noise ratio, occlusions, etc.) and the behavior of the fly (rigidity, leg injury, etc.) on a scale from 1 to 6 (where 1 is very good, 3 is satisfying, 6 is insufficient) for each 10 min recording session. We only retained sessions in which both criteria were at least at a satisfying quality level. Unless indicated otherwise, we analyzed trials where the fly was walking before stimulus onset. Thus, we did not retain data from flies with less than ten trials of walking before stimulation. We chose to do this for several reasons: (1) GCaMP6s decays very slowly. Even if the fly was moving *∼*2 s before stimulation, we still observed residual fluorescence signals, increasing the variability of changes upon stimulation. There were only very few instances where the animal was robustly resting for *>*2 s making the inverse analysis impossible. (2) We observed that control flies became aroused upon laser light stimulation. Thus, they may begin moving if they were resting prior to stimulation, indirectly driving DN activity and making it harder to discriminate between optogenetically-induced versus arousal-induced activity. We nevertheless provide data from flies that were resting before stimulation in **Supplementary File 1**, noting that the recruitment patterns, while not identical, are largely similar. DNp09 shows strong activation in the medial cervical connective (as for when the fly was walking prior to stimulation) and additional activation in lateral regions. The central neurons characteristic of aDN2 activation in animals that were previously walking are also active in animals that were previously resting. Additionally, we observe more widespread, weaker activation. DN signals upon MDN activation were slightly more spread out when the fly was resting prior to stimulation.

For experiments with headless animals, we excluded data from flies where one of the legs was visibly immobile after decapitation or when the abdomen was stuck to the spherical treadmill such that other movements became impossible.

### Behavioral data analysis

For analysis, we used custom Python code unless otherwise indicated. Code for behavioral data preprocessing can be found in the ‘twoppp’ Python package on GitHub previously used in ref. ^82^. Code for more detailed analysis can be found in the GitHub repository for this paper.

### Velocity computation

As a proxy for walking velocities, we tracked rotations of the spherical treadmill using Fictrac ^83^. Data from an IR camera placed in front of the fly was used for these measurements as described in ref. ^28^. Raw velocity traces acquired at 100 Hz were noisy and thus low pass filtered with a median filter (width = 5 = 0.05 s) and a Gaussian filter (*σ* = 10 = 0.1 s).

The velocity of the cylindrical treadmill was computed as follows. First, the wheel was detected in a camera on the left side of the fly using Hough circle detection. For each frame, we extracted a line profile along the surface of the wheel showing the dot pattern painted on its side. We then compared this line profile to the line profile of the previous frame to determine the most likely rotational shift. We converted this shift to a difference in wheel angle and then transformed this into a linear velocity in *mm/s* to make it comparable to quantification of spherical treadmill rotations. This image processing was prone to high frequency noise. Therefore, we filtered raw velocities with a Gaussian filter (*σ* = 20 = 0.2 s).

### 2D Pose estimation using SLEAP

We tracked nine keypoints from a camera on the right side of the fly: anal plate, ovipositor, most posterior stripe, neck, front leg coxa, front leg femur tibia joint, front leg tibia tarsus joint, mid leg tibia tarsus joint, hind leg tibia tarsus joint (see **Fig. 1d**) using SLEAP version 1.3.0^84^. All leg key-points were of right-side limbs on the fly.

### Behavior classification

We classified behaviors using an interpretable classifier based on heuristic thresholds of the walking velocity, limb motion energy (ME), and front leg height. For example, we classified forward and backward walking as having a forward velocity *>* 1 mm/s and *≤ −*1 mm/s, respectively. All parameters are shown in **Table 4**. If none of the conditions were fulfilled, we classified the behavior as undefined.

**Table 4:** Parameters used for behavior classification.

| Behavior | $v_{abs}$ (mm/s) | $v_{forw}$ (mm/s) | fr. leg height | ME front | ME mid | ME hind |
| --- | --- | --- | --- | --- | --- | --- |
| Rest | $\leq 0.75$ | n.a. | $> 0$ | $\leq 0.75$ | $\leq 0.75$ | $\leq 0.75$ |
| Forward walk | n.a. | $> 1$ | n.a | n.a | n.a | n.a |
| Backward walk | n.a. | $\leq -1$ | n.a | n.a | n.a | n.a |
| Anterior grooming 1 | $\leq 0.75$ | n.a. | $\leq 0$ | n.a. | $\leq 0.75$ | $\leq 0.75$ |
| Anterior grooming 2 | $\leq 0.75$ | n.a. | $> 0$ | $> 0.75$ | $\leq 0.75$ | $\leq 0.75$ |
| Posterior grooming | $\leq 0.75$ | n.a. | $> 0$ | $\leq 0.75$ | n.a. | $> 0.75$ |

Anterior grooming was composed of a logical ‘OR’ of two conditions: (1) The front leg was lifted up high, or (2) the front leg was moving with high motion energy. Front leg height was computed as the vertical distance between the front leg tibia-tarsus joint and the median position of the coxa. Pixel coordinates start from the top of the image. Thus, it is positive when the front leg is low (e.g., during resting) and negative when the front leg is high (e.g., during head grooming). Motion energy of the front, mid, and hind legs was computed based on the movements of the respective tibia-tarsus joint as follows: 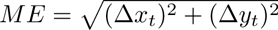, where Δ*_xt_* and Δ*_yt_* are the difference in *x* and *y* between two consecutive frames. We then computed the moving average of the motion energy within a 0.5 s (i.e., 50 samples) window to focus on longer time-scale changes in motion energy.

### Two-photon microscopy image analysis

We used custom Python code unless otherwise indicated. For all image analysis, the y-axis is dorsal-ventral along the fly’s body, and the x-axis is medial-lateral. Image and filter kernel sizes are specified as (y, x) in units of pixels. Code for two-photon data pre-processing can be found in the ‘twoppp’ Python package on GitHub previously used in ref. ^82^. Code for more detailed analysis can be found in the GitHub repository for this paper.

### Motion correction

Recordings from the thoracic cervical connective suffer from large inter-frame motion including large translations, as well as smaller, non-affine deformations. Contrary to motion correction procedures used before for similar data ^82^, here we made use of the high baseline fluorescence seen in Dfd*>*LexAop-GCaMP6s animals instead of relying on an additional, red color channel for motion correction. Thus, we performed motion correction directly on the green GCaMP channel. We compared the performance for data where a red channel was available and could only find negligible differences in ROI signals. Importantly, whether a neuron was encoding walking or resting was unchanged irrespective of whether we used the GCaMP channel or recordings from an additional red fluorescent protein.

We performed center of mass registration on every microscopy frame to compensate for large cervical connective translations. We then cropped the microscopy images (from 480×736 to 320×736 pixels). Then, we computed the motion field for each frame relative to one selected frame per fly using optic flow. We corrected the frames for this motion using bi-linear interpolation. The algorithm for optic flow motion correction was previously described in ref. ^51^. We only used the optic flow component to compute the motion fields and omitted the feature matching constraint. We regularized the gradient of the motion field to promote smoothness (*λ* = 800).

### ROI detection

For each pixel, we computed the standard deviation image across time for the entire recording. This gives a good proxy of whether a pixel belongs to a neuron—it has high standard deviation, because the neuron was sometimes active—or not. We used this image as a spatial map of the recording to inform ROI detection. Example standard deviation images are also used as the background image for **Fig. 2**c.

We applied principal component analysis on a subset of all pixels in the two-photon recording. We then projected the loadings of the first five PCs back into the image space. This gave us additional spatial maps integrating functional information to identify neurons. We then used a semi-automated procedure to detect ROIs: we performed peak detection in the standard deviation map. We visually inspected these peaks for correctness by looking at both the standard deviation map and the PCA maps. We manually added ROIs that the peak detection algorithm had missed, for example because the neuron was only weakly active. The functional PCA maps allowed us to discriminate between nearby neurons with dissimilar functions. They might show up as one big peak in the standard deviation map, but would clearly be assigned to different principal components. We were able to annotate between 50 and 80 ROIs for each fly. The number of visible neurons varies due to GCaMP6s expression levels, dissection quality, recording quality, and the behavioral activity level of the fly.

### Neural signal processing

We extracted fluorescence values for each annotated ROI by averaging all pixels within a rhomboid shape placed symmetrically over the ROI center (11 px high and 7 px wide). This gave us raw fluorescence traces across time for each neuron/ROI. We then low-pass filtered those raw fluorescence traces using a median filter (width= 3 =*∼* 0.185 s) and a Gaussian filter (*σ* = 3 =*∼* 0.185 s).

### Δ*F/F* computation

Because of variable expression levels among cells, GCaMP fluorescence is usually reported as a change in fluorescence relative to a baseline fluorescence. Here, we were mostly interested whether neurons were activated or not. To have a quantification that was comparable across neurons, we also normalised fluorescence of each neuron to its maximum level. Thus, we computed 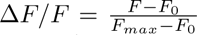 where *F* is the time-varying fluorescence of a neuron, *F*_0_ is its fluorescence baseline and *F_max_*is its maximum fluorescence. We computed *F_max_* as the 95 % quantile value of F across the entirety of the recording. In rare instances, neurons would get occluded, or slight glitches of the motion correction algorithm would result in some residual movement. Both of these make it challenging to estimate the minimum fluorescence. When the fly is resting, nearly all neurons are at their lowest levels (aside from several ^28^) and there is usually less movement of the nervous system. Thus, we computed *F*_0_ as a ‘resting baseline’ as follows. First, using our behavioral classifier we identified the onsets of prolonged resting (at least 75 % of 1 s after onset classified as resting and at least 1 s after the previous onset of resting) outside of optogenetic stimulation periods. For each neuron, we then computed the median fluorescence across repetitions aligned to resting onset. We then searched for the minimum value in time over the 2 s following rest onset. Taking the median across multiple instances of resting provided a more stable way to compute the baseline than by simply taking the minimum fluorescence. The normalization using *F*_0_ and *F_max_* provided a way to compare fluorescence across multiple neurons with similar units. Thus, whenever we report absolute Δ*F/F*, a value of 0 refers to neural activity during resting and 1 refers to the 95 % quantile of neural activity. When we report Δ*F/F* relative to pre-stimulus values **(Fig. 2b-f,i, Extended Data Fig. 2)**, then the unit of Δ*F/F* persists and a value of 0.5 means that the neuron has changed its activity level half as much as when it would go from a resting state to its 95 % quantile state.

### Video data processing

To process the raw fluorescence videos shown in the **supplementary videos 1-7** and **Fig. 2b**, we first low-pass filtered the data with the same temporal filters as for ROI signals (median filter width=3=*∼* 0.185 s, Gaussian filter *σ* = 3 =*∼* 0.185 s). Additionally, we applied spatial filters (median filter width=(3,3) px, Gaussian filter *σ* = (2, 2) px). We then applied the same Δ*F/F* computation method described above, but for each individual pixel instead of for individual ROIs. Thus, the units used in the videos are identical to the units used for ROI signals in **Fig. 2, Extended Data Fig. 1**.

### Synchronization of two-photon and camera data

We recorded two different data modalities at two different sampling frequencies: two-photon imaging data was recorded at approximately 16.23Hz, behavioral images from cameras were acquired at 100Hz. We synchronized these recordings using a trigger signal acquired at 30kHz. When it was necessary to analyse neural and behavioral data at the same sampling rate (e.g., **Supplementary Videos 1-7**), we down-sampled all measurements to the two-photon imaging frame rate by averaging all behavioral samples acquired during one two-photon frame. In the figures, we report data at its original sampling rate.

### Stimulus-triggered analysis of neural and behavioral data

We proceeded in the same way irrespective of whether the trigger was the onset of optogenetic stimulation **(Fig. 2**, **Fig. 4**, **Fig. 5, Extended Data Fig. 1, Extended Data Fig. 3)** or the onset of a natural (spontaneous, or puff-elicited) behavior **(Extended Data Fig. 2)**. To compute stimulus-triggered averages, we aligned all trials to the onset of stimulation and considered the times between 5 s prior to the stimulus onset and 5 s after stimulus offset. In **Fig. 2** we only considered trials where the fly was walking in the 1 s prior to stimulation (behavior classification applied to the mean of the 1 s pre-stimulus interval). We only considered flies with at least 10 trials of walking before stimulation. Behavioral responses in **Fig. 2a**, **Fig. 4b-g**, **Fig. 5f-g, Extended Data Fig. 1d-f, Extended Data Fig. 2a-c, Extended Data Fig. 3** show the average across all trials (including multiple animals) and the shaded area indicates the 95 % confidence interval of the mean across trials. When behavioral probabilities are shown, the fraction of trials that a certain behavior occurs at a specific time after stimulus onset is shown. Neural responses over time in **Fig. 2d, Extended Data Fig. 2a-c** show average responses across all trials of one animal. In order to visualise the change in neural activity upon stimulation, the mean of neural activity in the 1 s prior to stimulation is subtracted for each neuron. If the absolute value of the mean across trials for a given neuron at a given time point was less than the 95 % confidence interval of the mean, the data was masked with 0 (i.e., it is white in the plot). This procedure allowed us to reject noisy neurons with no consistent response across trials. Because we subtracted the baseline activity before stimulus onset, we also observed DNs that became less active upon optogenetic stimulation (neurons appearing blue). However, GCaMP6s fluorescence does not reliably reflect neural inhibition. Thus, we cannot claim that this reduced activation in some neurons is due to inhibition. Instead, because the flies were walking before stimulation onset, those neurons most likely encode walking and became less active when the fly stopped walking forward.

Individual neuron responses in **Fig. 2c, Extended Data Fig. 2a-c,f** show the maximum response of a single neuron/ROI. We detected the maximum response during the first half of the stimulus (2.5 s). We then computed the mean response of this neuron during 1 s centered around the time of its maximum response. If during at least half of that 1 s the mean was confidently different from 0 (i.e., *|mean| > CI*), we considered the neuron to be responsive, otherwise we masked the response to zero to reject noisy neurons with no consistent response across trials. **Fig. 2b** shows the same as **Fig. 2c**, but with this processing applied to pixels rather than individual neurons/ROIs. Contrary to previous ROI processing, pixels are not masked to 0 in case they are not responsive. **Fig. 2e** shows an overlay of **Fig. 2c** for multiple flies. Data from each of these flies were registered to one another by aligning the y-coordinates of the most dorsal and ventral neurons, as well as the x-coordinate of the most lateral neurons. **Fig. 2f** is a density visualization of **Fig. 2e**. To compute the density, we set the individual pixel values where a neuron was located to its response value and summed this across flies. We then applied a Gaussian filter (*σ* = 25 px, kernel normalized such that it has a value of 1 in the center to keep the units interpretable) and divided by the number of flies to create an ‘average fly’. **Extended Data Fig. 2d** was generated in the same manner.

### Statistical tests

**Fig. 2g-i** include a statistical analysis of neural responses. We quantified the number of activated neurons for each fly **(g)** as the neurons whose response value was positive (as in **Fig. 2c**). We quantified the fraction of activated neurons for each fly **(h)** by dividing the number of activated neurons by the number of neurons detected in the recording. In **(i)**, we quantified the summed Δ*F/F* as the sum of the response values of neurons that were positively activated (see red line in **Fig. 2d**). Here, we ignored neurons with negative response values because reductions in GCaMP fluorescence should not be interpreted as reflecting inhibition (see above). We used two-sided Mann-Whitney U tests (*scipy.stats.mannwhitneyu* ^85^) to statistically analyze these comparisons. Sample sizes and p-values are described in the figure legends. The Mann-Whitney U test is a ranked test. Thus, comparing three samples against three samples (e.g., aDN2 versus control), where all samples are at identical relative positions (i.e., ranks), will yield the same p-value, even if the absolute values are slightly different. This leads the p-values to be identical across **Fig. 2g-i** reflecting the conservative choice of a rank test that does not assume an underlying distribution.

**Fig. 4b-e**, **Fig. 5f,g, Extended Data Fig. 3** show statistical tests comparing the behavioral responses of intact and headless flies. **Fig. 4f,g**, **Fig. 5f,g, Extended Data Fig. 3** show statistical tests comparing the behavioral responses of headless experimental flies with headless control flies. In each case, we used two-sided Mann-Whitney U tests (*scipy.stats.mannwhitneyu* ^85^) to compare the average value within the first 2.5 s after stimulus onset. We averaged across technical replicates (trials) and only compare biological replicates (individual flies) using statistical tests. Exact p-values rounded to three digits are indicated in **Table 5**.

**Table 5:** Exact p-values for statistical tests in headless animal experiments.

| Figure | GAL4 | Variable | Comparison | N | P-value |
| --- | --- | --- | --- | --- | --- |
| Fig. 4b | DNp09 | forw. vel. | intact/headless | 5 | 0.006 |
| Fig. 4b | DNp09 | forw. prob. | intact/headless | 5 | 0.006 |
| Fig. 4c | aDN2 | forw. vel. | intact/headless | 5 | 0.018 |
| Fig. 4c | aDN2 | groom prob. | intact/headless | 5 | 0.006 |
| Fig. 4d | MDN | forw. vel. | intact/headless | 5 | 0.500 |
| Fig. 4d | MDN | back prob. | intact/headless | 5 | 0.265 |
| Fig. 4e | control | forw. vel. | intact/headless | 5 | 0.500 |
| Fig. 4e | control | rest prob. | intact/headless | 5 | 0.018 |
| Fig. 4f | DNp09 v. control | abd. contr. | headless/headless | 5 | 0.006 |
| Fig. 4g | aDN2 v. control | fr. leg appr. | headless/headless | 5 | 0.030 |
| Fig. 5f | DNb02 | turn vel. | intact/headless | 3/7 | 0.001 |
| Fig. 5f | control | turn vel. | intact/headless | 5 | 0.047 |
| Fig. 5f | DNb02 v. control | turn vel. | headless/headless | 7/5 | 0.313 |
| Fig. 5f | DNg14 | abd. dip | intact/headless | 9 | 0.144 |
| Fig. 5f | control | abd. dip | intact/headless | 5 | 0.072 |
| Fig. 5f | DNg14 v. control | abd. dip | headless/headless | 9/5 | 0.003 |
| Extended Data Fig. 3a | aDN1 | fr. leg motion | intact/headless | 5 | 0.002 |
| Extended Data Fig. 3a | control | fr. leg motion | intact/headless | 5 | 0.006 |
| Extended Data Fig. 3a | aDN1 v. control | fr. leg motion | headless/headless | 5 | 0.006 |
| Extended Data Fig. 3a | aDN1 | FeTi angle | intact/headless | 5 | 0.008 |
| Extended Data Fig. 3a | control | FeTi angle | intact/headless | 5 | 0.417 |
| Extended Data Fig. 3a | aDN1 v. control | FeTi angle | headless/headless | 5 | 0.148 |
| Extended Data Fig. 3a | aDN1 | fr. leg appr. | intact/headless | 5 | 0.149 |
| Extended Data Fig. 3a | control | fr. leg appr. | intact/headless | 5 | 0.072 |
| Extended Data Fig. 3a | aDN1 v. control | fr. leg appr. | headless/headless | 5 | 0.030 |
| Extended Data Fig. 3b | DNa01 | turn vel. | intact/headless | 6 | 0.003 |
| Extended Data Fig. 3b | control | turn vel. | intact/headless | 5 | 0.047 |
| Extended Data Fig. 3b | DNa01 v. control | turn vel. | headless/headless | 6/5 | 0.206 |
| Extended Data Fig. 3b | DNa01 | side vel. | intact/headless | 6 | 0.003 |
| Extended Data Fig. 3b | control | side vel. | intact/headless | 5 | 0.047 |
| Extended Data Fig. 3b | DNa01 v. control | side vel. | headless/headless | 6/5 | 0.392 |
| Extended Data Fig. 3c | DNa02 | turn vel. | intact/headless | 6 | 0.002 |
| Extended Data Fig. 3c | control | turn vel. | intact/headless | 5 | 0.047 |
| Extended Data Fig. 3c | DNa02 v. control | turn vel. | headless/headless | 6/5 | 0.158 |
| Extended Data Fig. 3c | DNa02 | side vel. | intact/headless | 6 | 0.003 |
| Extended Data Fig. 3c | control | side vel. | intact/headless | 5 | 0.047 |
| Extended Data Fig. 3c | DNa02 v. control | side vel. | headless/headless | 6/5 | 0.085 |
| Extended Data Fig. 3d | DNb02 | turn vel. | intact/headless | 3/7 | 0.001 |
| Extended Data Fig. 3d | control | turn vel. | intact/headless | 5 | 0.047 |
| Extended Data Fig. 3d | DNb02 v. control | turn vel. | headless/headless | 7/5 | 0.313 |
| Extended Data Fig. 3d | DNb02 | forw. vel. | intact/headless | 3/7 | 0.028 |
| Extended Data Fig. 3d | control | forw. vel. | intact/headless | 5 | 0.500 |
| Extended Data Fig. 3d | DNb02 v. control | forw. vel. | headless/headless | 7/5 | 0.313 |
| Extended Data Fig. 3e | DNg14 | abd. dip | intact/headless | 9 | 0.144 |
| Extended Data Fig. 3e | control | abd. dip | intact/headless | 5 | 0.072 |
| Extended Data Fig. 3e | DNg14 v. control | abd. dip | headless/headless | 9/5 | 0.003 |
| Extended Data Fig. 3f | Mute | ovi. ext. | intact/headless | 3 | 0.500 |
| Extended Data Fig. 3f | control | ovi. ext. | intact/headless | 5 | 0.338 |
| Extended Data Fig. 3f | Mute v. control | ovi. ext. | headless/headless | 3/5 | 0.117 |

**Extended Data Fig. 2a-c (right),e** show the Pearson correlation between neural responses to optogenetic stimulation and neural activity during natural (spontaneous or puff-elicited) behaviors. The significance of the correlation is measured as the probability that a random sample has a correlation coefficient as high as the one reported (*scipy.stats.pearsonr* ^85^). In all figures showing statistical tests, significance levels are indicated as follows: *** = p*<*0.001, ** = p*<*0.01, * = p*<*0.05, n.s. = p*≥*0.05.

### Brain connectome analysis

#### Loading connectome data

We used the female adult fly brain (FAFB) connectomics dataset^3^ from Codex (version hosted on Codex as of 3 August 2023, FlyWire materialization snapshot 630, https://codex.flywire.ai/api/ download) to generate all figures. We merged the “Neurons”, “Morphology Clusters”, “Connectivity Clusters”, “Classification”, “Cell Stats”, “Labels”, “Connections”, and “Connectivity Tags” tables. We then found DNs by filtering for the attribute *super class=descending*. We identified DNs with known, named (e.g., DNp09) genetic driver lines from Namiki et al. ^14^ by checking the ‘Cell Type’, ‘Hemibrain Type’ and ‘Community Labels’ attributes (in this priority) and using the following rules. Otherwise, we used the consensus cell type ^86^ (e.g. DNpe078). We semi-automatically assigned names using the following rules:

1. Special neurons: we manually labelled root IDs 720575940610236514, 720575940640331472, 720575940631082808, and 720575940616026939 as MDNs based on community labels from Salil Bidaye (consensus cell type DNpe078); root IDs 720575940616185531 and 720575940624319124 as aDN1 based on community labels from Katharina Eichler and Stefanie Hampel (consensus cell type DNge197); and root IDs 720575940624220925 and 720575940629806974 as aDN2 based on community labels from Katharina Eichler and Stefanie Hampel (consensus cell type DNge078). We verified visually that the shape of the neurons corresponded to published light-level microscopy images ^20,18^.
2. Otherwise, if both the *hemibrain type* attribute and the *cell type* attribute followed the Namiki format (“DN*{*1 lowercase letter*}{*2 digits*}*”, eg. “DNp16”), and they are identical, we used this as the cell name. If they are both in this format but are not identical, we marked this neuron for manual intervention.
3. Otherwise, if the *hemibrain type* attribute follows the Namiki format, we used this as the cell name. In addition, if the *hemibrain type* attribute follows the Namiki format, but the the *cell type* attribute has a different value following the consensus cell type format (“DN*{*at least 1 lowercase letter*}{*at least 1 digit*}*”, like ‘DNge198’), we marked the cell as requiring manual attention.
4. Otherwise, if the *cell type* attribute follows the Namiki format, we used this as the cell name.
5. Otherwise, if the *cell type* attribute follows the consensus cell type format, we used this as the cell name.
6. Otherwise, we marked the cell as requiring manual intervention.
7. Wherever manual intervention was required (mostly where the *hemibrain type* is the Namiki format, but the *cell type* is in the consensus cell type format), we manually assigned the consensus cell type. However, we assigned the Namiki type if there was no other DN in this Namiki cell type or if the cell type was still missing a pair of DNs ^14^.

Next, we stored the connectome as a graph using SciPy sparse matrix ^85^ and NetworkX Directed-Graph ^87^ representations. We identified DNs with somas in the GNG by checking the third letter of the consensus cell type to be ‘g’ (i.e., DNgeXXX) ^86^.

#### Analyzing connectivity

We only considered neurons with at least five synapses to be connected and computed the number of connected DNs based on this criterion **(Fig. 3**, **Fig. 5b,c,e, Fig. 6a-c, Extended Data Fig. 3)**. This is the same value as the default in Codex, the connectome data explorer provided by the FlyWire community ^42^. Analysis of connectivity across three brain hemispheres (2 brain halves from the FAFB dataset ^3^ and one from the hemibrain dataset ^88^) revealed that connections ‘stronger than ten synapses or 1.1 % of the target’s inputs have a greater than 90 % change to be preserved” ^86^. We visualized all DNs connected to a given DN **(Fig. 3a,b**, **Fig. 5d, Extended Data Fig. 3)** using the neuromancer interface and manually colored neurons depending on whether they are in the GNG or not.

Neurotransmitter identification was available from the connectome dataset based on classification of individual synapses with an average accuracy of 87 % ^43^. Here, we report neurotransmitter identity for a given presynaptic-postsynaptic connection. To define neurotransmitter identity for a given presynaptic-postsynaptic pair, we asserted that the neurotransmitter type would be unique using a majority vote rule. This was chosen as a tradeoff between harmonizing neurotransmitters for a neuron (especially GABA, acetylcholine and glutamate ^89^) and avoiding the propagation of classification errors.

#### DN network visualisations and DN hierarchy

We used the networkx library ^87^ to plot networks of DNs in **Fig. 3c,d**, **Fig. 5e, Extended Data Fig. 3**. Again, we considered neurons to be connected if they had at least five synapses. In the circular plots we show summed connectivity of multiple DNs. For example, the network for DNp09 in **Fig. 3c** shows only one green circle in the center representing two DNp09 neurons. All connections shown as arrows are the sum of those two neurons. DNs are considered excitatory if they have the neurotransmitter acetylcholine and inhibitory if they have the neurotransmitter GABA. Whether glutamate is excitatory or inhibitory is unclear—this depends on the receptor subtype ^75^ which is unknown in most cases. To emphasize this, we highlight glutamatergic network edges in a different color (pink).

In figure **Fig. 3e** we show the cumulative distribution of the number of DNs reachable within up to N synapses. Statistics on DN connectivity across multiple synapses were computed using matrix multiplication with the *numpy* library on the adjacency matrix of the network. Colored lines represent a DN network traversal starting at specific command-like DNs. The black trace represents the median of all neurons. Only a maximum of approximately 800 DNs can be reached because the others have maximally one DN input. In **Fig. 5b,c** we sorted DNs by the number of monosynaptic connections they make to other DNs. In **Fig. 5b**, the same sorting is applied to show the number of connected GNG DNs (orange).

#### Fitting network models to connectivity degree distribution

In **Fig. 6a**, we generated a shuffled network of the same size by keeping the number of neurons constant and keeping the number of connections constant. Then, we randomly shuffled (i.e., reassigned) those connections. Here, we only considered the binary measure of whether a neuron was connected or not (number of synapses *>*5) and not its synaptic weight. We then fit a power-law or an exponential to the connectivity degree distribution using the *scipy.optimize* ^85^ library. Histograms of the degree distributions for all four distributions are shown in **Fig. 6a** using constant bin widths of 5 neurons. The quality of the fits are quantified using linear mean squared error (*R*^2^).

#### Detection of DN clusters

We applied the Louvain method ^50^ with resolution parameter *γ* = 1 to detect clusters. Briefly, the Louvain method is a greedy algorithm that maximizes modularity (i.e., the relative density of connections within clusters compared to between clusters). To simplify the network during the optimization, we do not consider the directionality of connections between neurons. If there is reciprocal connectivity between neurons, we add up the number of synapses (positive if excitatory, negative if inhibitory; here glutamate is considered inhibitory and neuromodulators are disregarded for the sake of simplicity). The Louvain method finds different local optima of cluster assignments due to its stochastic initialization and greedy nature. Therefore, we ran the algorithm 100 times. Based on the outcomes of these 100 runs, we define a co-clustering matrix: The matrix has the same size as the connectivity matrix (*number of DNs × number of DNs*). Each entry represents how often two DNs end up in the same cluster. This matrix assigns each pair of DNs a probability to be in the same cluster.

We then applied hierarchical clustering to this matrix (using the ‘ward’ optimization method from the *scipy.cluster.hierarchy* library ^85^) to get the final sorting of DNs shown in **Fig. 6b**. Using this meta-clustering, we could be sure that the sorting of DNs we find through clustering is not a local optimum and that it is reproducible. We then used the final sorting to detect the clusters shown in grey in **Fig. 6b** as follows: We started from one side of the sorted DNs and sequentially grew the cluster. If the next DN was in the same Louvain clusters at least 25 % of the time, we assigned it to the same cluster as the previous DN. If not, we started a new cluster with this DN and kept testing subsequent DNs as to whether they fulfill the criteria for this new cluster. Finally, we only kept clusters that had at least 10 neurons. This yielded 12 clusters (grey squares). We applied the same meta-clustering technique to a shuffled network (same number of DNs, same number of connections, same number of synapses, but shuffled connections). On this shuffled network, we found 34 clusters of much smaller size **(Fig. 6c)**, hinting at a better clustering in our network than in a shuffled control (*modularity* = 0.27 for the original network and *modularity* = 0.12 for the shuffled network). The number of synapses is shown as positive values (red) if it is excitatory and as negative values (blue) if it is inhibitory.

We then analyzed the connectivity within and between clusters. To do this, we accumulated the number of synapses between two clusters (positive for excitatory, negative for inhibitory). In order to be able to compare this quantity between clusters of different sizes, we divided this number of synapses by the number of DNs in the cluster that receives the synaptic connections. This quantity is visualized in **Fig. 6d** for the original DN-DN network clusters and **Fig. 6e** for the shuffled network as the ‘normalized number of synapses’. If positive (red), then connections from one cluster to another are predominantly excitatory. If negative (blue) connections are predominantly inhibitory.

#### Statistical comparison of original versus shuffled DN-DN clusters

As detailed above, we applied the Louvain algorithm 100 times to increase the robustness of clustering. We computed statistics on the clustering of this dataset (mean and standard deviation) specifically on metrics including the size and number of clusters. We then compared these distributions with those for the shuffled graph using one-sided t-tests (*scipy.stats.ttest ind* ^85^). The resulting statistics are a conservative quantification of the difference between the original network and the shuffled control, as each data point is taken independently. When performing the hierarchical clustering across 100 iterations, the large clusters from the biological network are preserved whereas the random associations of the shuffled network become incoherent. In practice, the difference in cluster sizes reported statistically underestimates the difference between the resulting matrices shown in **Fig. 6b,c**. The 100 iterations result from random seed initialization, on the condition that the algorithm converges. We restarted it whenever the convergence criteria was not reached within 3 s. Indeed we observed empirically that when the algorithm would not converge in 3 s it would not do so for at least 30 min and was, therefore, terminated.

#### Identifying DNs to test predictions

Based on the cell type data associated with each neuron in FAFB (see above), we were able to find many DNs from refs. ^14,15,18^ in the connectome database. We then checked which of them have either a very high number of synaptic connections to other DNs or a very low number. We then filtered for lines where a clean split-GAL4 line was available. Additionally, we focused on lines whose major projections in the VNC were outside of the wing neuropil, because we removed the wings in our experimental paradigm and thus might not be able to see optogenetically-induced behaviors. This left us with ten additional DNs to test our predictions. DNp01 (giant fiber) activation was reported to trigger take-off in intact and headless flies ^46,47^ so we did not repeat those experiments. This left us with nine lines to test. The source and exact genotypes of those fly lines are reported in **Table 2**. We then performed experiments with those nine lines. Because intact control flies become aroused by laser illumination, but not headless control animals, to avoid false positives we only analyzed DN lines that either had a known optogenetic behavior in intact flies (i.e., aDN1, DNa01, DNa02), or that had a clear phenotype in headless flies (i.e., DNb02, DNg14, Mute). Thus, we excluded Web, DNp24, and DNg30 as they did not fulfill either of these criteria and only analyzed the remaining six driver lines in **Fig. 5** and **Extended Data Fig. 3**.

#### Analyzing DN-DN connectivity in the VNC

We used the neuprint website to interact with the male adult nerve cord (MANC) connectome dataset ^9,65^. There, we searched neurons based on their name (MDN, DNp09, etc.) and checked whether there were any DNs among the neurons that are postsynaptic. We found all neurons that we used from ref. ^14^ (i.e., DNp09, DNa01, etc.), and MDN. We were not able to find aDN2, aDN1, Mute and Web.

#### Analyzing VNC targets of DN clusters

We used data shown in Cheong et al. ^9^ Figure 3 - Supplement 2 to define whether a DN known from Namiki et al. ^14^ was projecting to a particular VNC neuropil. Briefly, a DN is considered as projecting to a given neuropil if at least 5 % of its presynaptic sites are in that region. We manually found the MDNs in the MANC dataset and determined the regions they connect to using the same criterion. To generate **Fig. 6f**, for each cluster, we accumulated the number of known DNs that project to a given VNC region. We then divided this by the number of known DNs to obtain the fraction of known DNs within a cluster that project to a given region. The number of unknown DNs per cluster is also shown next to the plot. The raw data of associations between DNs and VNC neuropils is shown in **Supplementary Data File 2**.

#### Analyzing behaviors associated with DN clusters

We examined the literature ^12,19,18,9,51,52,53,20,17,90,29,13,54,10^ to identify behaviors associated with DNs and grouped them into broad categories (anterior movements, take-off, landing, walking, flight). This literature summary is available in the **Supporting Information File 2**. Of the 34 DN types annotated, we found conflicting evidence for only two: DNg11 is reported to elicit foreleg rubbing ^19^ while targeting mostly flight-related neuropils ^9^; DNa08 targets flight power control circuits ^9^ but has been reported to be involved in courtship under the name aSP22^22^. In **Fig. 6g** we assigned DNg11 to ‘anterior’ and DNa08 to ‘flight’. We accumulated the number of known DNs that are associated with a given behavior for each cluster. We then divided by the number of known DNs in the respective cluster to get a fraction of DNs within a cluster that have a known behavior. The number of unknown DNs per cluster is also shown next to the plot. The raw data of associations between DNs and behaviors is shown in **Supplementary Data File 2**.

#### Analyzing brain input neuropils for each DN cluster

We used data from FAFB to identify the brain input neuropils for each DN cluster based on the neuropil annotation for each DN-DN synapse. Thus, localization information is given by the position of each synaptic connection and not the cell body of the presynaptic partner. This allows us to account for local processing and modularity of neurons. The acronyms of brain regions are detailed in **Table 6**, with ‘L’ and ‘R’ standing for the left and right brain hemispheres respectively. Results are reported as the fraction of synapses made in a neuropil out of all the post-synaptic connections made by DNs of a given cluster.

**Table 6:** Acronyms for the brain neuropils used in Fig. 6h based on ref. ^3^.

| Acronym | Neuropil | Region |
| --- | --- | --- |
| AL | antennal lobe | Antennal Lobe |
| AME | accessory medulla | Optic Lobe |
| AMMC | antennal mechanosensory and motor center | Periesophageal Neuropils |
| AOTU | anterior optic tubercle | Ventrolateral Neuropils |
| ATL | antler | Inferior Neuropils |
| AVLP | anterior VLP (ventrolateral protocerebrum) | Ventrolateral Neuropils |
| BU | bulb | Lateral Complex |
| CAN | cantle | Periesophageal Neuropils |
| CRE | crepine | Inferior Neuropils |
| EPA | epaulette | Ventromedial Neuropils |
| FB | fanshaped body | Central Complex |
| FLA | flange | Periesophageal Neuropils |
| GA | gall | Lateral Complex |
| GNG | gnathal ganglia | Gnathal Ganglia |
| GOR | gorget | Ventromedial Neuropils |
| IB | inferior bridge | Inferior Neuropils |
| ICL | inferior clamp | Inferior Neuropils |
| IPS | inferior posterior slope | Ventromedial Neuropils |
| LAL | lateral accessory lobe | Lateral Complex |
| LH | lateral horn | Lateral Horn |
| LOP | lobula plate | Optic Lobe |
| LO | lobula | Optic Lobe |
| MB_CA | pedunculus | Mushroom Body |
| MB_ML | vertical lobe | Mushroom Body |
| MB_PED | medial lobe | Mushroom Body |
| MB_VL | calyx | Mushroom Body |
| NO | noduli | Central Complex |
| OCG | ocellar ganglion | Ocelli |
| PB | fanshaped body | Central Complex |
| PLP | posteriorlateral protocerebrum | Ventrolateral Neuropils |
| PRW | prow | Periesophageal Neuropils |
| PVLP | posterior VLP (ventrolateral protocerebrum) | Ventrolateral Neuropils |
| SAD | saddle | Periesophageal Neuropils |
| SCL | superior clamp | Inferior Neuropils |
| SIP | superior intermediate protocerebrum | Superior Neuropils |
| SLP | superior lateral protocerebrum | Superior Neuropils |
| SMP | superior medial protocerebrum | Superior Neuropils |
| SPS | superior posterior slope | Ventromedial Neuropils |
| UNASGD | unassigned |  |
| VES | vest | Ventromedial Neuropils |
| WED | wedge | Ventrolateral Neuropils |

## Data availability

Data are available at: https://dataverse.harvard.edu/dataverse/dn_networks This repository includes processed data required to reproduce the figures for each fly. Raw two-photon imaging data and behavioral videos (*∼* 1 TB) are available upon request.

## Code availability

Analysis code is available at: https://github.com/NeLy-EPFL/dn_networks

## Supporting Information Files

Supporting Information File 1 - Individual fly neural and behavioral responses to optogenetic stimulation (Ref. **Fig. 2**)

**Part 1: Flies were walking prior to stimulation.** Each page is for a single genotype. Each row shows individual fly neural and behavioral results. The number of trials is indicated. The analyses and plots are as in **Fig. 2a,c,d**. The bottom row summarizes data across all flies of the genotype as in **Fig. 2a,e,f**. Flies were only used if they had at least 10 trials of forward walking before stimulation onset.

**Part 2: Flies were resting prior to stimulation.** Same as for **Part 1** except that flies were resting before to stimulation onset. Flies were only used if they had at least 10 trials of resting before stimulation onset.

Link to Supporting Information File 1

Supporting Information File 2 - DN cluster analysis

(Ref. **Fig. 6**)

**Sheet 1: DN cluster behaviors.** A list showing which DNs are in which particular cluster.

**Sheet 2: DN cluster VNC projections.** VNC projections for known DNs. Aside from MDN, these data were obtained from ^9^.

**Sheet 3: Investigated DN clusters.** A subset of sheet 1 showing the cluster associations for DNs investigated using optogenetics in this study: DNp09, aDN2, MDN, aDN1, DNa01, DNa02, DNb02, DNg14 and Mute.

**Sheet 4: Equivalence DN names and root id.** List of DNs, providing the known names for the root ids used in FAFB ^3^.

**Sheet 5: DN literature aggregation** Reported behavioral phenotypes for DNs, including citations.

**Sheet 6: Connectivity statistics.** Connectivity statistics in the DN-DN network for reported neurons.

Link to Supporting Information File 2

## Supplementary Videos

**Supplementary Video 1: DNp09-driven behavior and trial-averaged GNG-DN population activity** (top) Stimulus-triggered average of neural activity upon DNp09 optogenetic stimulation. Video shows Δ*F/F* processed GCaMP6s fluorescence. (bottom) Four instances of forward walking for one animal (fly in **Fig. 2b-d**) upon DNp09 stimulation. Red circle indicates laser stimulation.

Link to Supplementary Video 1

**Supplementary Video 2: aDN2-driven behavior and trial-averaged GNG-DN population activity** (top) Stimulus-triggered average of neural activity upon aDN2 optogenetic stimulation. Video shows Δ*F/F* processed GCaMP6s fluorescence. (bottom) Four instances of anterior grooming for one animal (fly in **Fig. 2b-d**) upon aDN2 stimulation. Red circle indicates laser stimulation.

Link to Supplementary Video 2

**Supplementary Video 3: MDN-driven behavior and trial-averaged GNG-DN population activity** (top) Stimulus-triggered average of neural activity upon MDN optogenetic stimulation. Video shows Δ*F/F* processed GCaMP6s fluorescence. (bottom) Four instances of backward walking for one animal (fly in **Fig. 2b-d**) upon MDN stimulation. Red circle indicates laser stimulation.

Link to Supplementary Video 3

**Supplementary Video 4: Control for light-driven behavior and trial-averaged GNG-DN population activity** (top) Stimulus-triggered average of neural activity upon optogenetic stimulation of a fly without a GAL4 driver. Video shows Δ*F/F* processed GCaMP6s fluorescence. (bottom) Four instances of behavior for one animal (fly in **Fig. 2b-d**) upon light exposure. Red circle indicates laser exposure.

Link to Supplementary Video 4

**Supplementary Video 5: Comparing GNG-DN population activity for DNp09-driven versus spontaneous forward walking.** Forward walking (left) driven by optogenetic stimulation of DNp09 versus (right) spontaneously generated in the same animal (fly in **Extended Data Fig. 2a**). (top) Stimulus-triggered average of neural activity. Video shows Δ*F/F* processed GCaMP6s fluorescence. (bottom) Four instances of behavior time-locked to stimulation or forward walking onset. Red circle indicates laser stimulation. White circle indicates spontaneous forward walking detection.

Link to Supplementary Video 5

**Supplementary Video 6: Comparing GNG-DN population activity for aDN2-driven versus puff-induced anterior grooming.**Anterior grooming (left) driven by optogenetic stimulation of DNp09 versus (right) elicited by vapor-puff stimulation in the same animal (fly in **Extended Data Fig. 2b**). (top) Stimulus-triggered average of neural activity. Video shows Δ*F/F* processed GCaMP6s fluorescence. (bottom) Four instances of behavior time-locked to stimulation or anterior grooming onset. Red circle indicates laser stimulation. White circle indicates vapor puff stimulation. Link to Supplementary Video 6

**Supplementary Video 7: Comparing GNG-DN population activity for MDN-driven versus spontaneous backward walking.** Backward walking (left) driven by optogenetic stimulation of MDN versus (right) spontaneously generated in the same animal on a cylindrical treadmill (fly in **Extended Data Fig. 2c**). (top) Stimulus-triggered average of neural activity. Video shows Δ*F/F* processed GCaMP6s fluorescence. (bottom) Four instances of behavior time-locked to stimulation or backward walking onset. Red circle indicates laser stimulation. White circle indicates spontaneous backward walking onset.

Link to Supplementary Video 7

**Supplementary Video 8: DNp09-driven behavioral responses of animals that are intact, headless, or headless without ground contact.** Responses to optogenetic stimulation of DNp09 for three flies (one animal per column). The same animal is studied intact on the spherical treadmill (top), headless on the spherical treadmill (middle), and headless while hanging without ground contact (bottom). Red circles indicate optogenetic laser stimulation.

Link to Supplementary Video 8

**Supplementary Video 9: aDN2-driven behavioral responses of animals that are intact, headless, or headless without ground contact.** Responses to optogenetic stimulation of aDN2 for three flies (one animal per column). The same animal is studied intact on the spherical treadmill (top), headless on the spherical treadmill (middle), and headless while hanging without ground contact (bottom). Red circles indicate optogenetic laser stimulation.

Link to Supplementary Video 9

**Supplementary Video 10: MDN-driven behavioral responses of animals that are intact, headless, or headless without ground contact.** Responses to optogenetic stimulation of MDN for three flies (one animal per column). The same animal is studied intact on the spherical treadmill (top), headless on the spherical treadmill (middle), and headless while hanging without ground contact (bottom). Red circles indicate optogenetic laser stimulation.

Link to Supplementary Video 10

**Supplementary Video 11: Control animal behavioral responses to laser illumination of animals that are intact, headless, or headless without ground contact.** Responses to laser illumination of animals without a GAL4 driver for three flies (one animal per column). The same animal is studied intact on the spherical treadmill (top), headless on the spherical treadmill (middle), and headless while hanging without ground contact (bottom). Red circles indicate laser illumination.

Link to Supplementary Video 11

**Supplementary Video 12: DNb02-driven behavioral responses of animals that are intact, headless, or headless without ground contact.** Responses to optogenetic stimulation of DNb02 for three flies (one animal per column). Animals are studied intact on the spherical treadmill (top), headless on the spherical treadmill (middle), or headless while hanging without ground contact (bottom). Red circles indicate optogenetic laser stimulation.

Link to Supplementary Video 12

**Supplementary Video 13: DNa01-driven behavioral responses of animals that are intact, headless, or headless without ground contact.** Responses to optogenetic stimulation of DNa01 for three flies (one animal per column). Animals are studied intact on the spherical treadmill (top), headless on the spherical treadmill (middle), or headless while hanging without ground contact (bottom). Red circles indicate optogenetic laser stimulation.

Link to Supplementary Video 13

**Supplementary Video 14: DNa02-driven behavioral responses of animals that are intact, headless, or headless without ground contact.** Responses to optogenetic stimulation of DNa02 for three flies (one animal per column). Animals are studied intact on the spherical treadmill (top), headless on the spherical treadmill (middle), or headless while hanging without ground contact (bottom). Red circles indicate optogenetic laser stimulation.

Link to Supplementary Video 14

**Supplementary Video 15: aDN1-driven behavioral responses of animals that are intact, headless, or headless without ground contact.** Responses to optogenetic stimulation of aDN1 for six flies (each animal identity is indicated). Animals are studied intact on the spherical treadmill (top), headless on the spherical treadmill (middle), or headless while hanging without ground contact (bottom). Red circles indicate optogenetic laser stimulation.

Link to Supplementary Video 15

**Supplementary Video 16: DNg14-driven behavioral responses of animals that are intact, headless, or headless without ground contact.** Responses to optogenetic stimulation of DNg14 for three flies (one animal per column). Animals are studied intact on the spherical treadmill (top), headless on the spherical treadmill (middle), or headless while hanging without ground contact (bottom). Red circles indicate optogenetic laser stimulation.

Link to Supplementary Video 16

**Supplementary Video 17: Mute-driven behavioral responses of animals that are intact, headless, or headless without ground contact.** Responses to optogenetic stimulation of Mute for six flies (each animal identity is indicated). Animals are studied intact on the spherical treadmill (top), headless on the spherical treadmill (middle), or headless while hanging without ground contact (bottom). Red circles indicate optogenetic laser stimulation.

Link to Supplementary Video 17

## Acknowledgments

We thank Jasper Phelps and Stefanie Boy-Röttger for help with confocal dissections and staining. We thank Ĺea Goffinet for help with the initial identification of DNs in the female adult fly brain connectome. We thank Daniel Morales for generating transgenic fly lines. We thank Julie Simpson, Orkun Akin, Salil Bidaye, and Brian McCabe for sharing transgenic fly lines. We thank Katharina Eichler, Stefanie Hampel, and Salil Bidaye for annotating DNs in the female adult fly brain connectome. We thank the members of the Neuroengineering lab for helpful discussions and comments on the manuscript. JB acknowledges support from a Boehringer Ingelheim Fonds PhD stipend. FH acknowledges support from a Boehringer Ingelheim Fonds PhD stipend. SW acknowledges support from a Boehringer Ingelheim Fonds PhD stipend. PR acknowledges support from an SNSF Project Grant (175667) and an SNSF Eccellenza Grant (181239).

## Author Contributions

J.B. - Conceptualization, Methodology, Software, Validation, Formal Analysis, Investigation, Data Curation, Validation, Writing – Original Draft Preparation, Writing – Review & Editing, Visualization.

Technical contributions: **Fig. 1** visualisation; **Fig. 2** experiments, analysis, visualisation; **Fig. 3** some visualisation; **Fig. 4** experiments, analysis, visualisation; **Fig. 5** some experiments, some data analysis, some visualisation.

F.H. - Conceptualization, Methodology, Software, Validation, Formal Analysis, Investigation, Data Curation, Validation, Writing – Review & Editing, Visualization.

Technical contributions: **Fig. 3** most connectomics data analysis, most visualisation; **Fig. 5** most experiments, most data analysis, some visualisation; **Fig. 6** connectomics modelling, visualisation.

S.W. - Conceptualization, Methodology, Software, Validation, Formal Analysis, Investigation, Data Curation, Validation, Writing – Review & Editing.

Technical contributions: **Fig. 3**, **Fig. 5** some connectomics data analysis

P.R. - Conceptualization, Methodology, Resources, Writing – Original Draft Preparation, Writing - Review & Editing, Supervision, Project Administration, Funding Acquisition.

## Ethical compliance

All experiments were performed in compliance with relevant national (Switzerland) and institutional (EPFL) ethical regulations.

---

**Extended Data Fig. 1:**
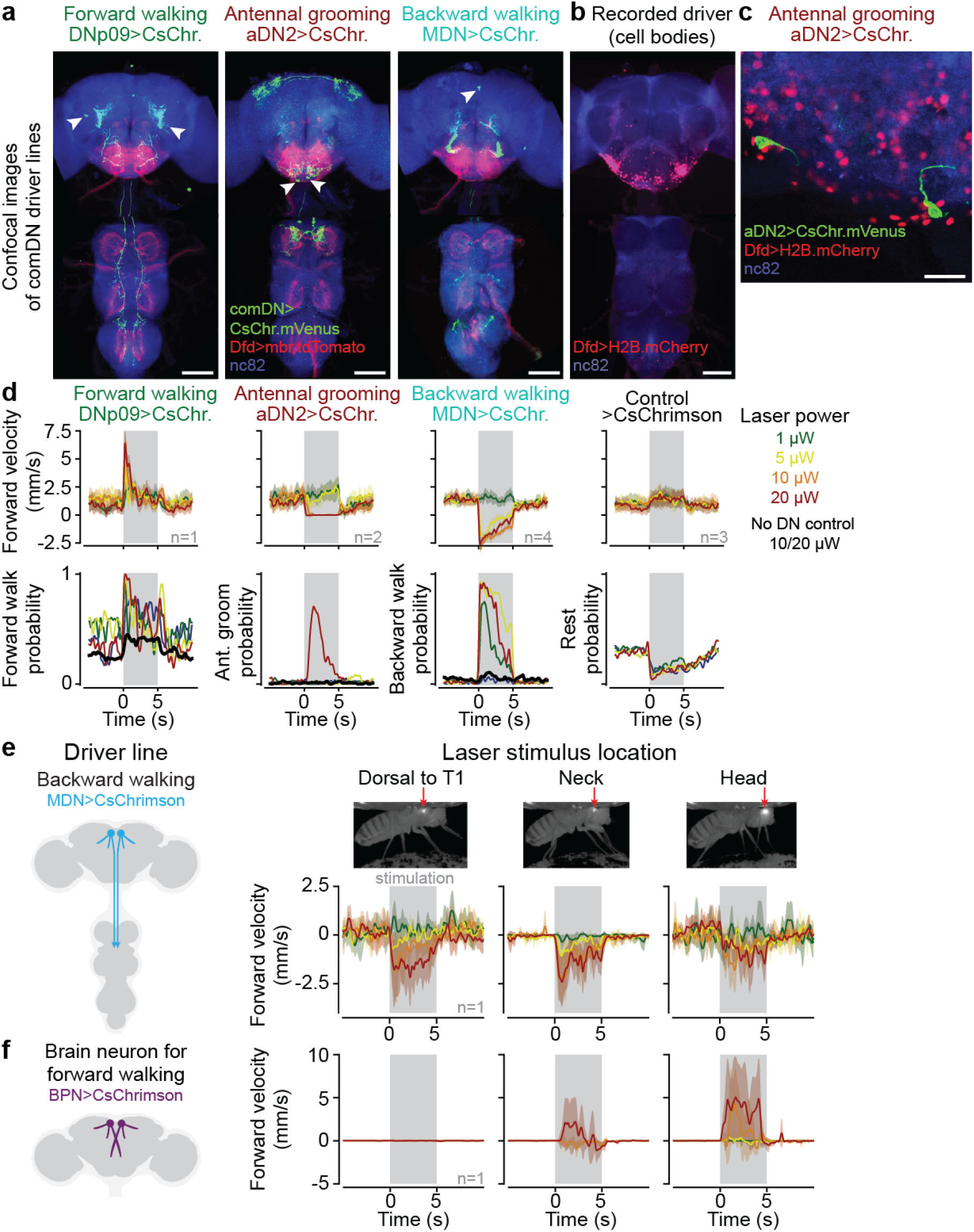
DN driver lines and optogenetic stimulation strategy. **(a)** Z-projected confocal images of the brain (top) and VNC (bottom) show the expression of UAS-CsChrimson.mVenus (green) in command-like DNs, membrane-bound tdTomato in the Dfd driver line (red), and neuropil (‘nc82’, blue). The location of command-like DN cell bodies is indicated (white arrowheads). Scalebars are 100 *µ*m. **(b)** Z-projected confocal image of Dfd driver line expression of soma-targeted mCherry. Only brain neurons in the GNG are labeled. Scalebar is 100 *µ*m. **(c)** Confocal image of the posterior GNG with Dfd driver line expression of soma-targeted mCherry and aDN2 expression of UAS-CsChrimson.mVenus (green). The two GNG-DNs in the aDN2 driver line are not targeted by the Dfd driver line. Scalebar is 20 *µ*m. **(d)** Behavioral responses to optogenetic stimulation of the neck connective at different laser intensities for DNp09 (left; 1 fly, 19 trials per condition), aDN2 (left-middle; 2 flies, 40 trials per condition), MDN (right-middle; 4 flies, 50 trials per condition), and no DN control (right; 3 flies, 60 trials per condition) animals. Flies reliably (i) walk forward upon DNp09 stimulation for stimuli *>* 5 *µ*W, (ii) groom upon aDN2 stimulation only for the highest stimulation power (20 *µ*W) but rest at 10 *µ*W, and (iii) backward walk upon MDN stimulation for stimuli *>* 5 *µ*W. For all stimulation intensities, control flies walk more and rest less. Thus, we selected 10 *µ*W as our default laser stimulation power and 20 *µ*W for aDN2 stimulation specifically. **(e)** MDN stimulation with focused laser light elicits backward walking when shining light at the anterior dorsal thorax (left, as in Fig. 4 and Fig. 5), at the neck (middle, as in Fig. 2), but less reliably at the head (right). N=6 stimulation trials per condition for one fly. **(f)** Stimulation of a brain-specific neuron (‘Bolt protocerebral neurons’ or BPN) known to drive forward walking ^17^ with focused laser light elicits forward walking when shining light on the head (right), but not on the thorax (left). Laser light focused on the neck (middle) can only elicit weak forward walking at 20 *µ*W. N=5 stimulation trials per condition for one fly. All traces in this figure show mean *±* 95 % confidence interval of the mean.

**Extended Data Fig. 2:**
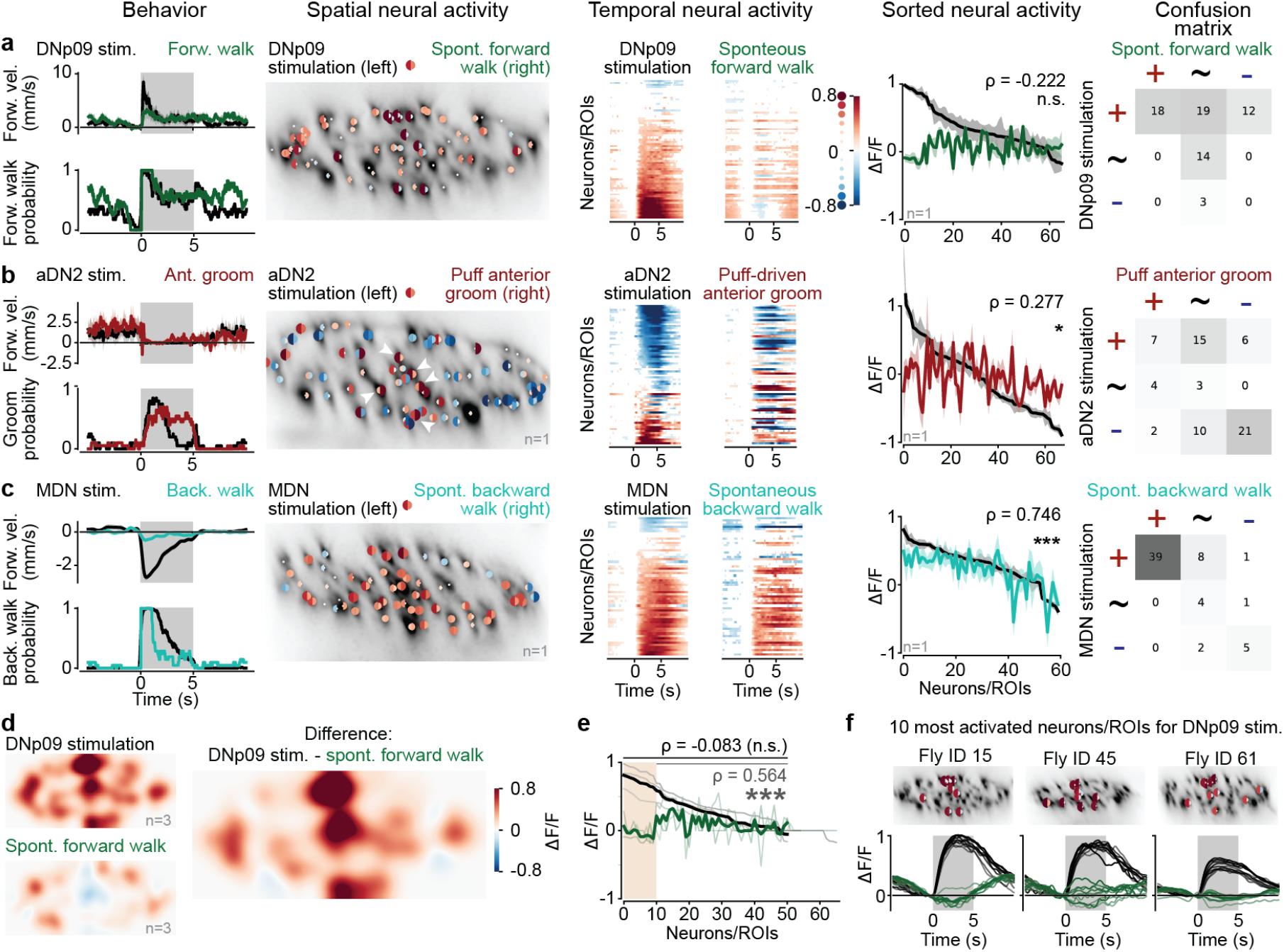
Comparison of GNG-DN population neural activity during optogenetic stimulation versus corresponding natural behaviors. **(a-c)** For **(a)** DNp09 and forward walking, **(b)** aDN2 and anterior grooming, or **(c)** MDN and backward walking: **(left)** behavioral responses to optogenetic stimulation of command-like DNs (black) versus natural occurrences of the behavior in question (color); **(middle left)** single neuron/ROI responses (analyzed as in Fig. 2c). Here the left half-circle reflects the response to optogenetic activation and the right half-circle the activity during natural behavior; **(middle)** single neuron average responses as in Fig. 2d; **(middle right)** Comparing the activity of individual neurons between optogenetic stimulation (black) and natural behavior (color). Neurons/ROIs are sorted by the magnitude of response to optogenetic activation. Shaded areas indicate 95% confidence interval of the mean across trials. Pearson correlation between optogenetic and spontaneous response and significance of test against null-hypothesis (the two variables are uncorrelated, see Methods) are shown; **(right)** Confusion matrix comparing the number of active neurons/ROIs that were more active (+), similar (*∼*), or less active (-) upon optogenetic stimulation versus during natural behavior. **(a)** DNp09: for one fly n=23 optogenetic stimulation trials (not forward walking before stimulus) and 28 instances of spontaneous forward walking in which the fly was not walking forward for at least 1 s and then walking forward for at least 1 s (correlation: *ρ* = *−*0.022*, p* = 0.356). **(b)** aDN2: for one fly, n=20 optogenetic stimulation trials (pre-stimulus behavior not restricted) and 16 instances of anterior grooming elicited by a 5 s humidified air puff (correlation: *ρ* = 0.277*, p* = 0.022). Indicated are central neurons/ROIs with strong activation during aDN2 stimulation of the neck cervical connective as in Fig. 2f. **(c)** MDN: for one fly, n=80 optogenetic stimulation trials (pre-stimulus behavior not restricted) and 21 instances of spontaneous backward walking on a cylindrical treadmill in which the fly was not walking backward for 1 s and then walked backward for at least 1 s (correlation: *ρ* = 0.746*, p <* 0.001). **(d)** Density visualisation (as in Fig. 2f) of neural responses to DNp09 stimulation and spontaneous forward walking across three animals. The difference in responses is primarily localized to the central but not lateral regions of the connective. To maximize comparability, only trials where the fly was not walking forward before stimulus onset were selected. **(e)** Same plot as in **a, middle right** but for three animals with DNp09 stimulation and forward walking. Indicated are the correlation values when including (*ρ* = *−*0.083*, p* = 0.564) or excluding (*ρ* = 0.564*, p <* 0.001) the ten neurons most activated by optogenetic stimulation (orange region). **(f)** The locations of ten neurons indicated in **e** within the connective of three flies (top) and their single neuron responses to optogenetic stimulation (bottom, black traces) or during natural backward walking (bottom, green traces).

**Extended Data Fig. 3:**
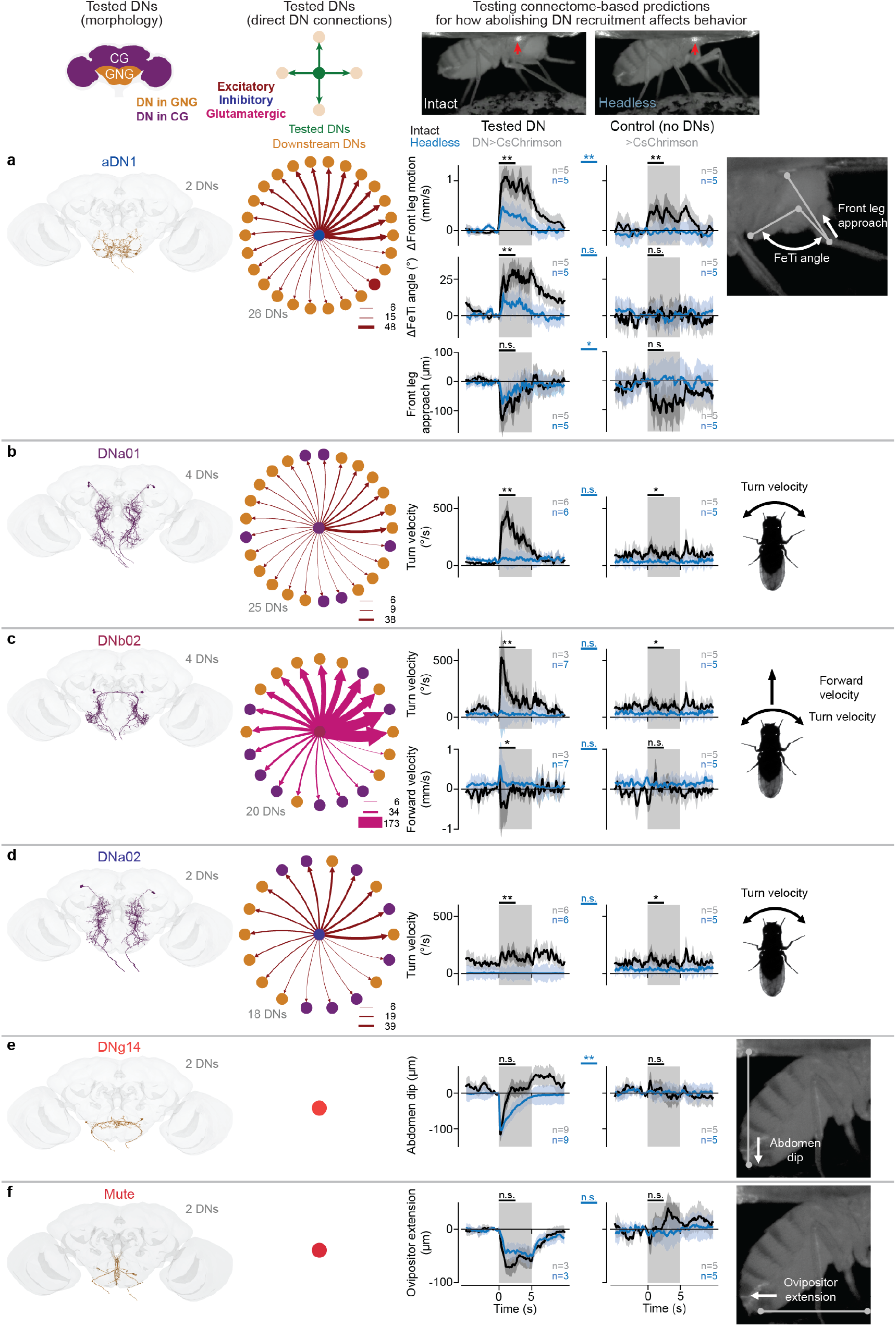
Testing connectome-based predictions of DN-driven behavioral flexibility and its dependence upon downstream DNs. **(a-f) (first column)** The morphology of tested DNs in the adult female brain connectome. DNs are color-coded based on their somata localization within the cerebral ganglia (purple) or gnathal ganglia (orange). The number of DNs is indicated. **(second column)** A network schematic of direct connections from tested to downstream DNs. Edge widths reflect the number of synapses and is consistent across plots. Edge colors denote excitatory (red), inhibitory (blue), or glutamatergic (pink) which can be excitatory or inhibitory depending on receptor type ^75^. **(third column)** Quantitative analyses of optogenetically-driven behaviors and movements in intact (black traces) and headless animals (blue traces). The number of flies for each condition are indicated. Each fly is optogenetically stimulated ten times. Thus, the average and 95 % confidence interval of the mean for a total of *n ∗* 10 trials is shown. **(fourth column)** Identical behavioral analysis for control flies without DN opsin expression. Note that controls for different parameters include the same five animals. Mann-Whitney U tests comparing the trial mean of intact and headless animals (black bars, above each plot) and comparing headless experimental with headless control flies (blue, in between experimental and control plots) are shown (*** means p*<*0.001, ** means p*<*0.01, * means p*<*0.05, n.s. means p*≥*0.05; for exact p-values see Methods). **(fifth column)** An illustration of the behavioral parameter(s) being quantified. **(a)** aDN1 has monosynaptic connections to 26 other DNs and triggers grooming in intact animals. By contrast, headless animals produce mostly uncoordinated front leg movements. These occur more slowly at a lower frequency (top) with a smaller change in femur-tibia angle (middle). The ‘front leg approach’ to the head—the change in Euclidean distance between the neck and tibia-tarsus joint relative to 1 s before stimulus onset—is similar between intact and headless animals (bottom). **(b)** DNa01 has monosynaptic connections to 25 other DNs and triggers in place turning. This is quantified as an increase in turn velocity. This behavior is lost in headless animals. **(c)** DNb02 has monosynaptic connections to 20 other DNs and weakly triggers turning. This is quantified as an increase in turning velocity (top), a phenotype that is lost in headless animals. Instead, a flexion of the front legs can be observed in headless animals. This is quantified as a short spike in forward velocity (bottom). These data partially overlap with those in **Fig. 5d-g**). **(d)** DNa02 has monosynaptic connections to 18 other DNs and weakly triggers turning. This is quantified as an increase in turning velocity. This behavior is lost in headless animals. **(e)** DNg14 has no monosynaptic connections to other DNs and triggers abdominal dipping and vibration in both intact and headless animals. This movement is quantified as a change in the vertical position of the anal plate relative to 1 s before stimulus onset. These are the same data as in **Fig. 5d-f**). **(f)** The DN ‘Mute’ has no monosynaptic connections to other DNs and triggers ovipositor extension in both intact and headless animals. This movement is quantified as a change in the horizontal position of the ovipositor relative to the 1 s before stimulus onset.

