## Supplementary Information File 1 for "Networks of descending neurons transform command-like signals into population-based behavioral control"

### Part 1:
### Individual fly neural and behavioral responses to optogenetic stimulation. Flies were **walking** prior to stimulation. (ref. Figure 2)

Braun, et al. 2023

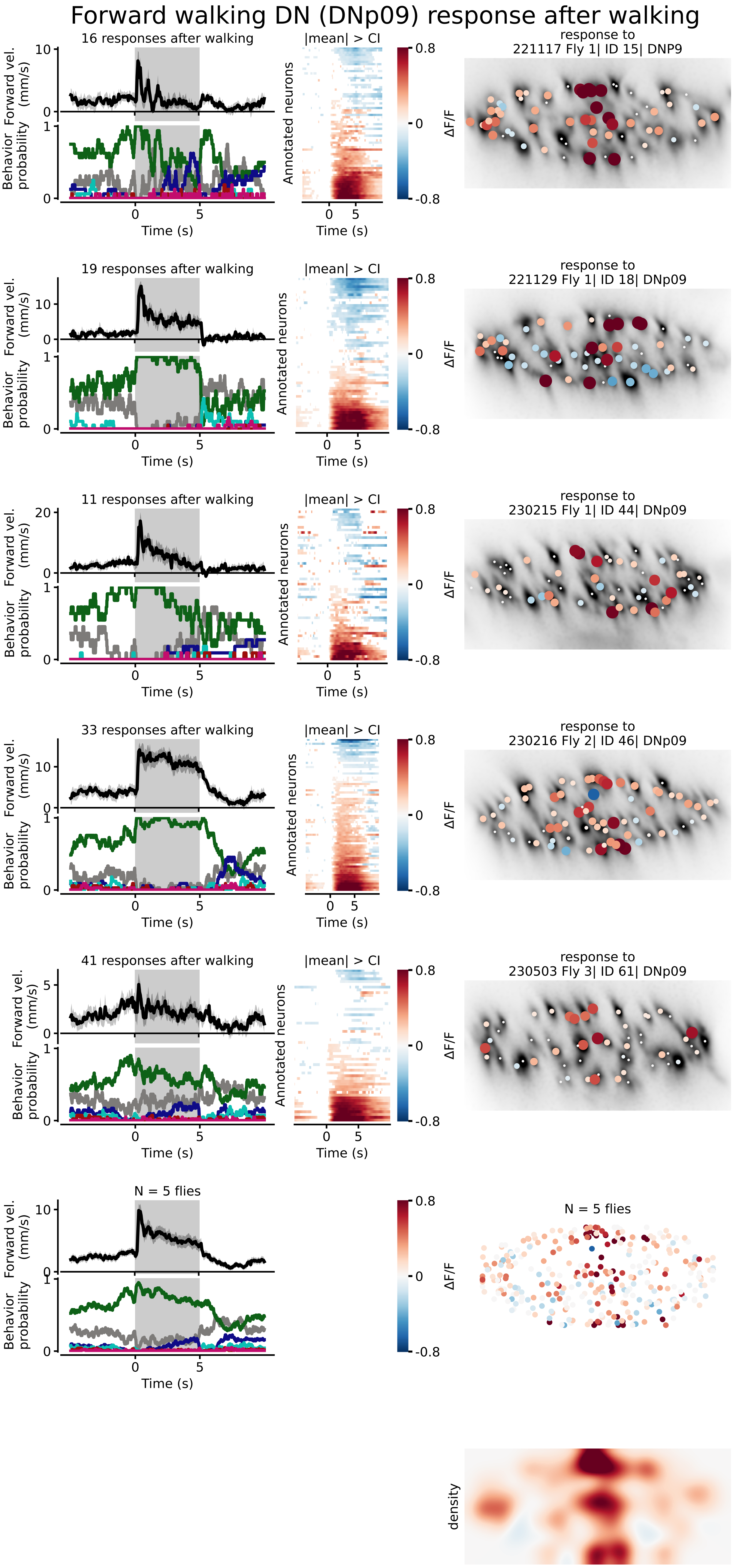
Forward walking DN (DNp09) response after walking
16 responses after walking
19 responses after walking
11 responses after walking
33 responses after walking
41 responses after walking
N = 5 flies
Forward vel. (mm/s)
Behavior probability
Time (s)
|mean| > CI
Annotated neurons
ΔF/F
density
response to 221117 Fly 1| ID 15| DNP9
response to 221129 Fly 1| ID 18| DNp09
response to 230215 Fly 1| ID 44| DNp09
response to 230216 Fly 2| ID 46| DNp09
response to 230503 Fly 3| ID 61| DNp09
N = 5 flies

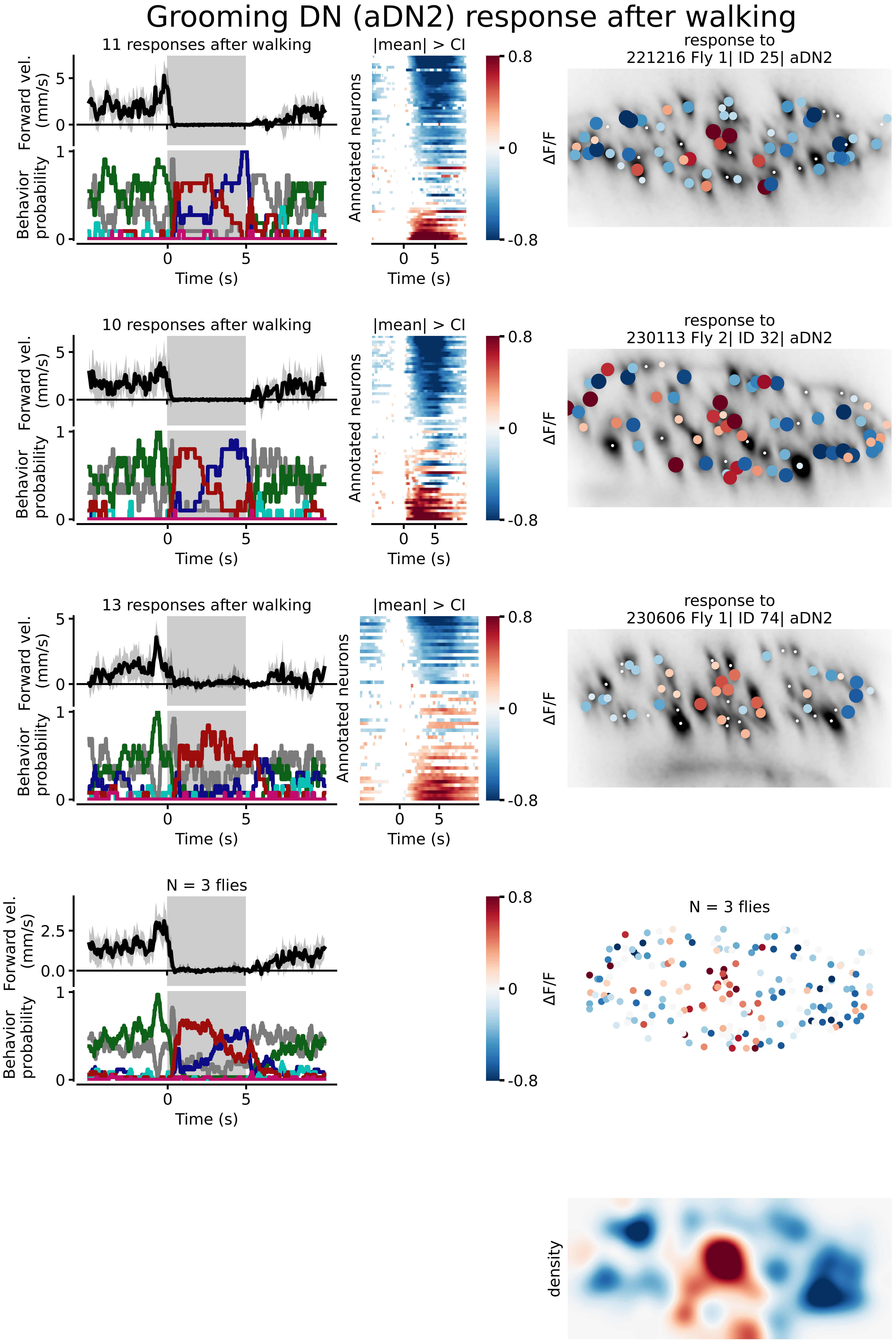
Grooming DN (aDN2) response after walking
11 responses after walking
10 responses after walking
13 responses after walking
N = 3 flies
Forward vel. (mm/s)
Behavior probability
Time (s)
|mean| > CI
Annotated neurons
ΔF/F
0.8
0
-0.8
response to
221216 Fly 1| ID 25| aDN2
response to
230113 Fly 2| ID 32| aDN2
response to
230606 Fly 1| ID 74| aDN2
N = 3 flies
density

### Backward walking DN (MDN) response after walking

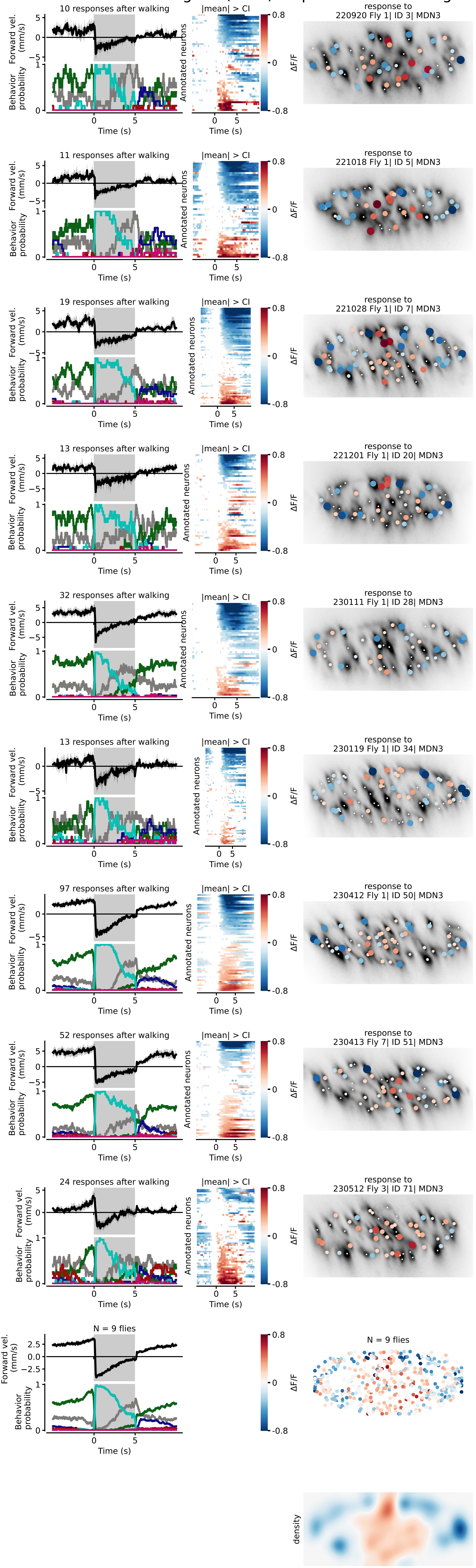

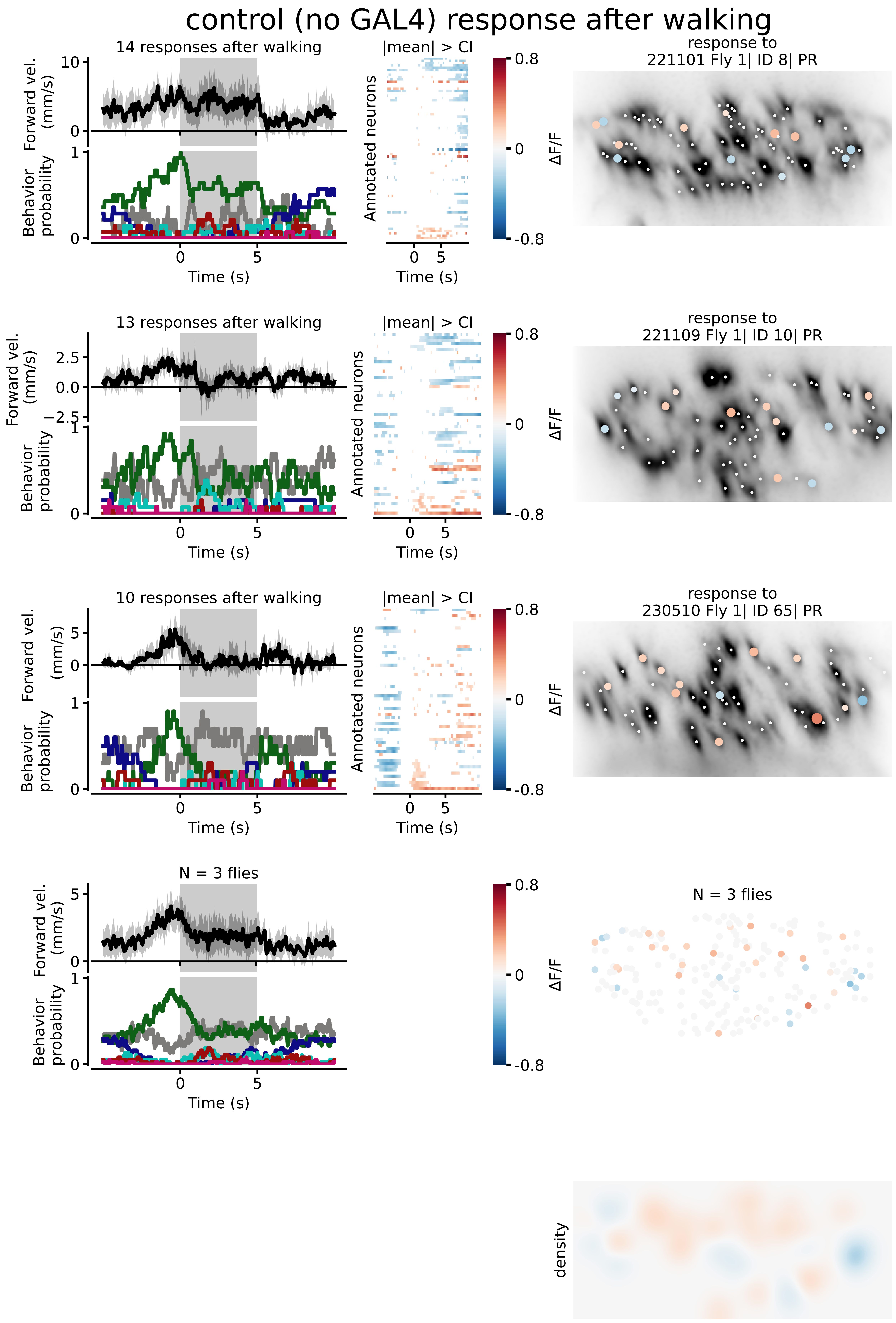

control (no GAL4) response after walking
14 responses after walking
Forward vel. (mm/s)
Behavior probability
Time (s)
|mean| > CI
Annotated neurons
response to 221101 Fly 1| ID 8| PR
ΔF/F
13 responses after walking
response to 221109 Fly 1| ID 10| PR
10 responses after walking
response to 230510 Fly 1| ID 65| PR
N = 3 flies
density

### Part 2:
### Individual fly neural and behavioral responses to optogenetic stimulation. Flies were **resting** prior to stimulation. (ref. Figure 2)

Braun, et al. 2023

### Forward walking DN (DNp09) response after resting

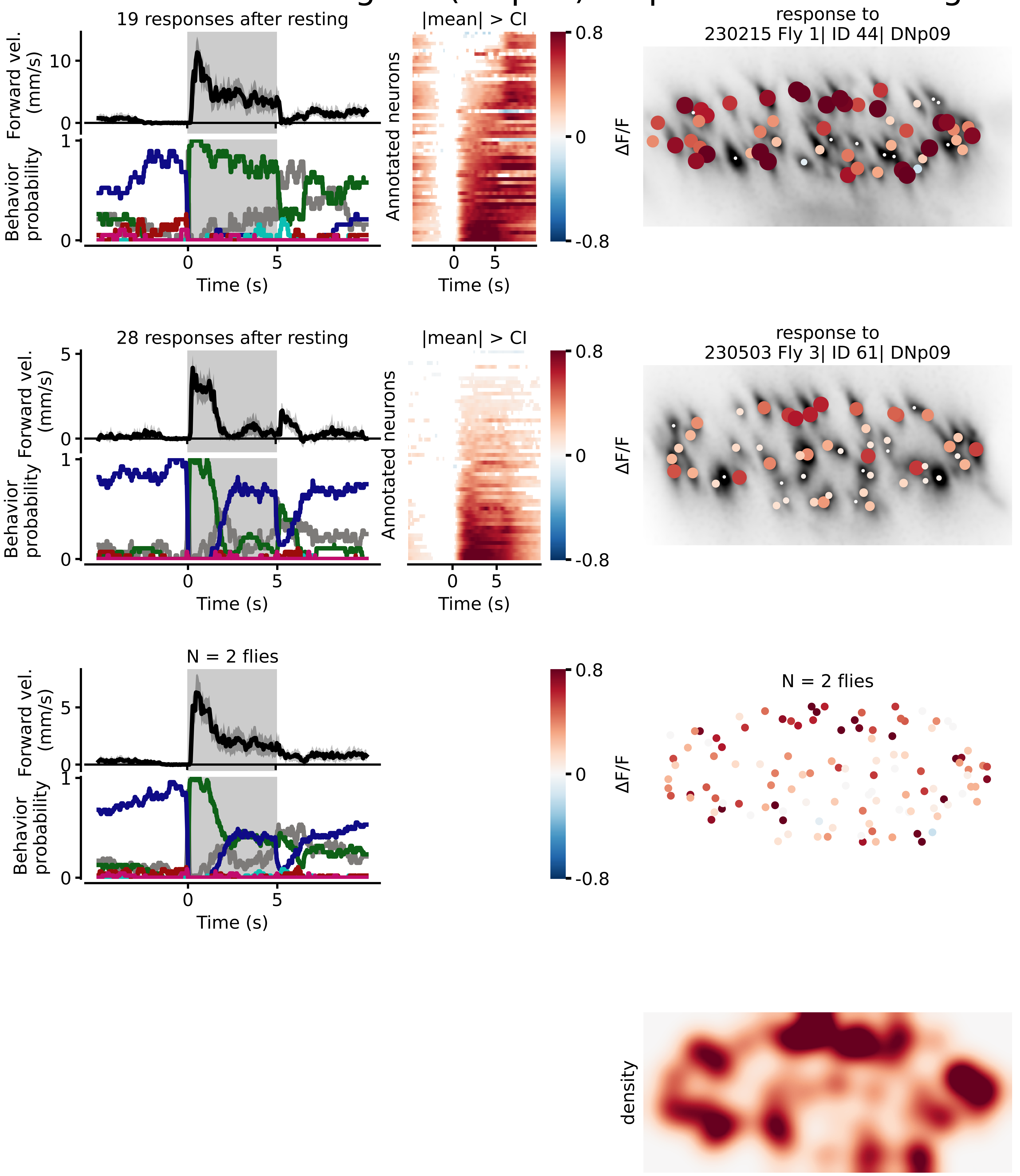

### Grooming DN (aDN2) response after resting

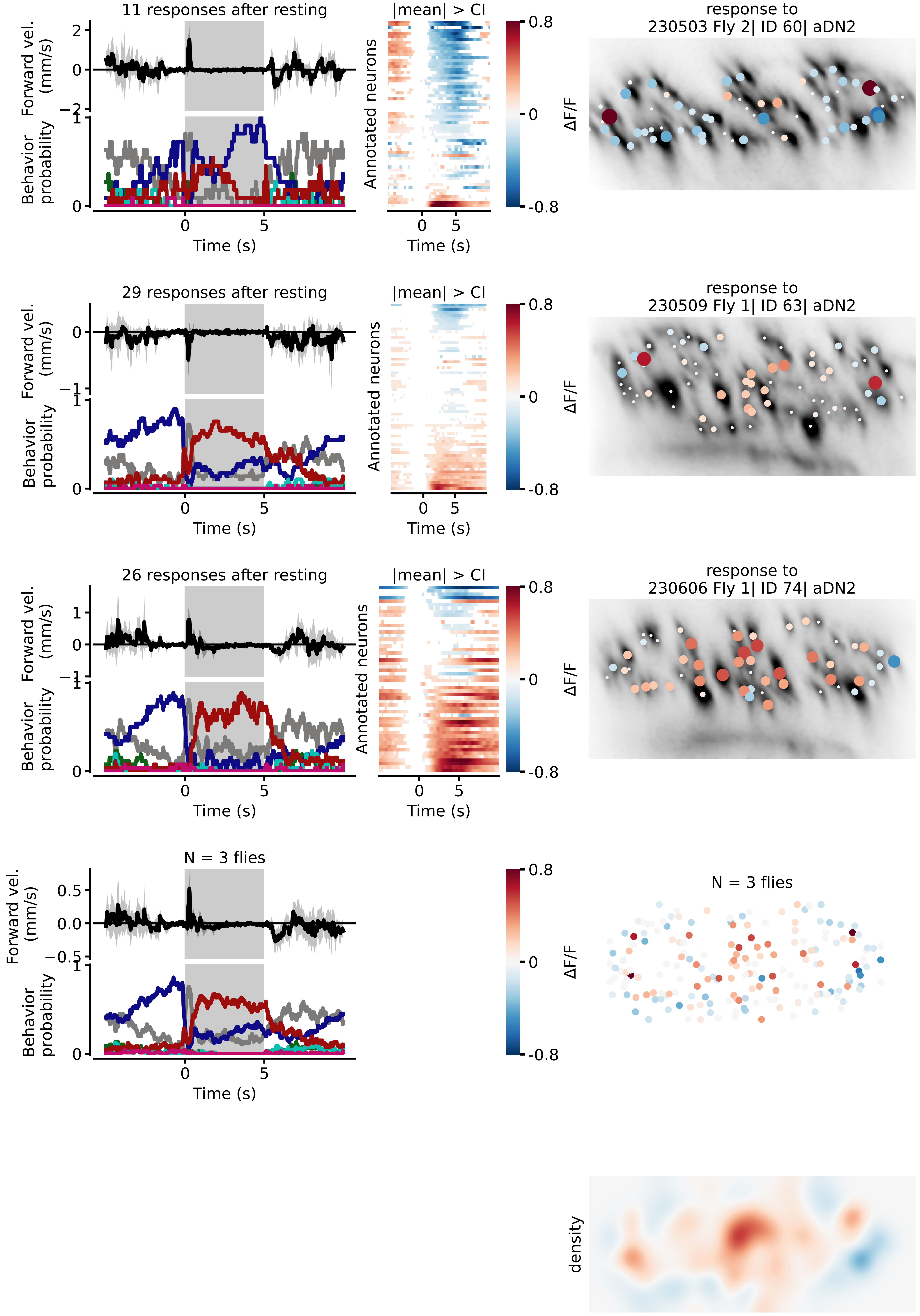

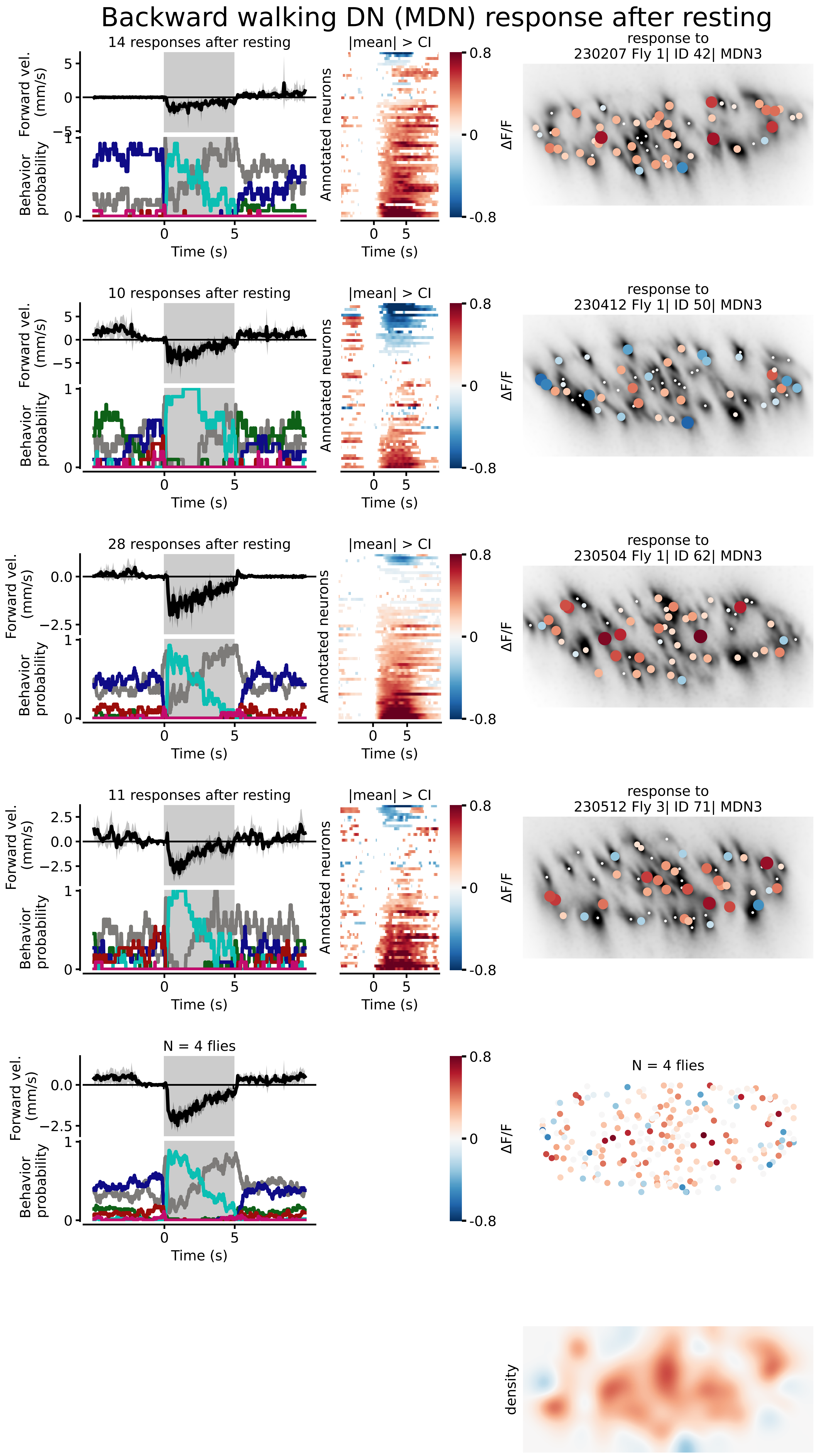
Backward walking DN (MDN) response after resting
14 responses after resting
10 responses after resting
28 responses after resting
11 responses after resting
N = 4 flies
Forward vel. (mm/s)
Behavior probability
Time (s)
|mean| > CI
Annotated neurons
ΔF/F
response to 230207 Fly 1| ID 42| MDN3
response to 230412 Fly 1| ID 50| MDN3
response to 230504 Fly 1| ID 62| MDN3
response to 230512 Fly 3| ID 71| MDN3
N = 4 flies
density

### control (no GAL4) response after resting

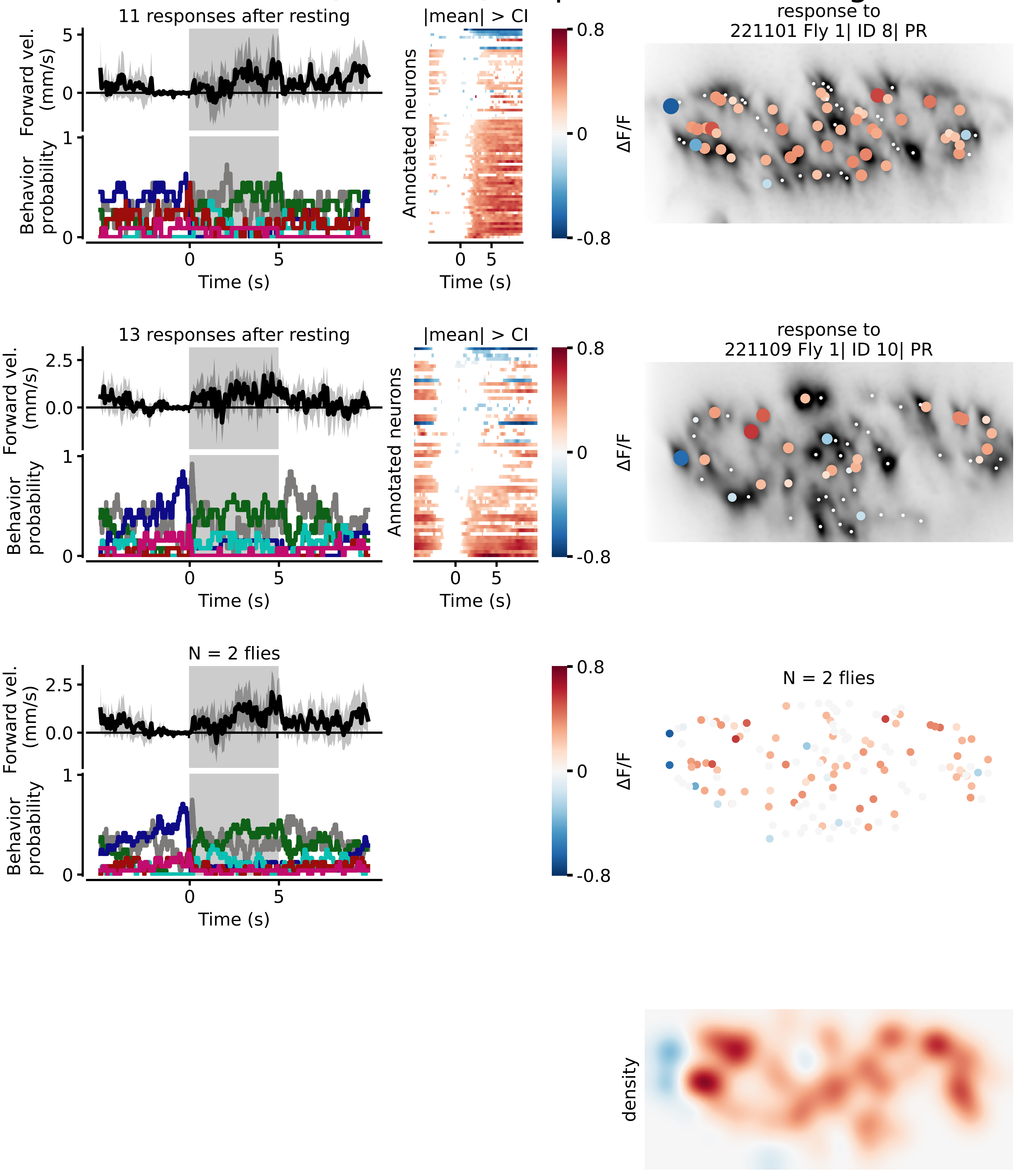
